# Lateral gene transfer introduced the microbial anaerobiosis-related gene *rquA* into early animals

**DOI:** 10.64898/2026.04.08.717022

**Authors:** Sofia Paraskevopoulou, Mara Vizitiu, Nikolaj Brask, Paige E. Carr, Julie Boisard, Humberto Itriago, Raquel Pereira, Iana V. Kim, Tuğba N. Atalay, Lily N. Boercker, Karla I. Aguilera Campos, Laura Flandrin, Disa Stephensen, Cai S. Westergren, Fabien Pierrel, Arnau Sebé-Pedrós, Jennifer N. Shepherd, April L. Horton, Sally P. Leys, Courtney W. Stairs

**Author notes:** Corresponding author: Courtney Stairs.

## Abstract

Lateral gene transfer (LGT) enables rapid metabolic innovation in microbes, but its evolutionary importance in animals remains debated. Among metabolic traits with major ecological consequences, adaptations to low-oxygen conditions often involve modifications of mitochondrial electron transport and the quinones that mediate electron flow. Rhodoquinone-based anaerobic metabolism occurs in several eukaryotic lineages, yet the evolutionary routes by which animals acquired this capability are poorly understood. Here we show that freshwater sponges possess a rhodoquinone biosynthesis gene, *rquA*, previously restricted to microbial lineages, which was acquired by lateral gene transfer and functionally integrated into sponge metabolism. Heterologous expression of *rquA* from the model freshwater sponge *Ephydatia muelleri* enables rhodoquinone production in yeast, consistent with functional conservation. In *E. muelleri*, the *rquA* gene is upregulated under hypoxia and rhodoquinone is detectable across all lifestages, however, it is most abundant in early development in the pluiripotent ‘gemmules’. Using comparative genomics, we find that the presence of *rquA* in freshwater sponges correlates with loss of key genes of the ubiquinone biosynthesis pathway, suggesting these animals cannot synthesize ubiquinone *de novo* and we show that *E. muelleri* can convert exogenous ubiquinone to rhodoquinone. Rhodoquinone levels were significantly higher in *rquA*-encoding freshwater sponges compared to marine sponges that were sampled from natural environments. This study reveals that an early animal lineage acquired a microbial metabolism-related gene via lateral gene transfer during or before the transition to freshwater environments, enabling rhodoquinone utilization and potentially enhancing tolerance to oxygen fluctuations. Thereby, demonstrating how LGT shapes energy metabolism even in multicellular organisms.

## Main

Lateral gene transfer (LGT), *i.e.,* the movement of genetic material across species boundaries, is central in microbial evolution, enabling rapid metabolic innovations and ecological diversification^1–4^. In animals, however, its evolutionary role has long been debated mainly due to constraints imposed by early germline segregation^5–8^. Growing evidence documents multiple LGT events from bacteria and fungi to animal lineages^9–12^, indicating that foreign gene acquisition is not restricted to unicellular life and may have contributed to metabolic and ecological diversification in metazoans.

Among traits potentially shaped by LGT, adaptations in metabolism have immense consequences, as they determine how organisms conserve energy^3,13^. The mitochondrial electron transport chain (ETC) of most aerobic eukaryotes consists of four large complexes important for energy conservation via Complex V (CV). Electrons from NADH or succinate enter the ETC via CI and CII, respectively, and are transferred to the benzoquinone ubiquinone (UQ)^13^ yielding ubiquinol (UQH_2_). Electrons from UQH_2_ are transferred through CIII, cytochrome c, and CIV and are ultimately deposited onto oxygen. Reactions catalyzed by CI, CIII and CIV are coupled to the translocation of protons from the matrix to the intermembrane space of the mitochondrion, yielding a proton gradient that fuels ATP biosynthesis via CV.

Under oxygen-limited conditions, some unicellular eukaryotes (protists) and animals express a truncated ETC that instead utilizes the benzoquinone rhodoquinone (RQ)^3,13,14^ (Fig. 1A). In this ETC, CI couples NADH oxidation and proton translocation to RQ reduction, however, the RQH_2_ is reoxidized by CII functioning in reverse. The lower reduction potential of RQ relative to UQ makes fumarate reduction more favourable allowing fumarate, and not oxygen, to serve as the terminal electron acceptor. This modified respiratory chain consisting of CI and CII is sufficient to maintain a proton gradient for ATP biosynthesis via CV in low oxygen (hypoxic) or anoxic environments and has evolved independently in several lineages.

**Fig. 1:**
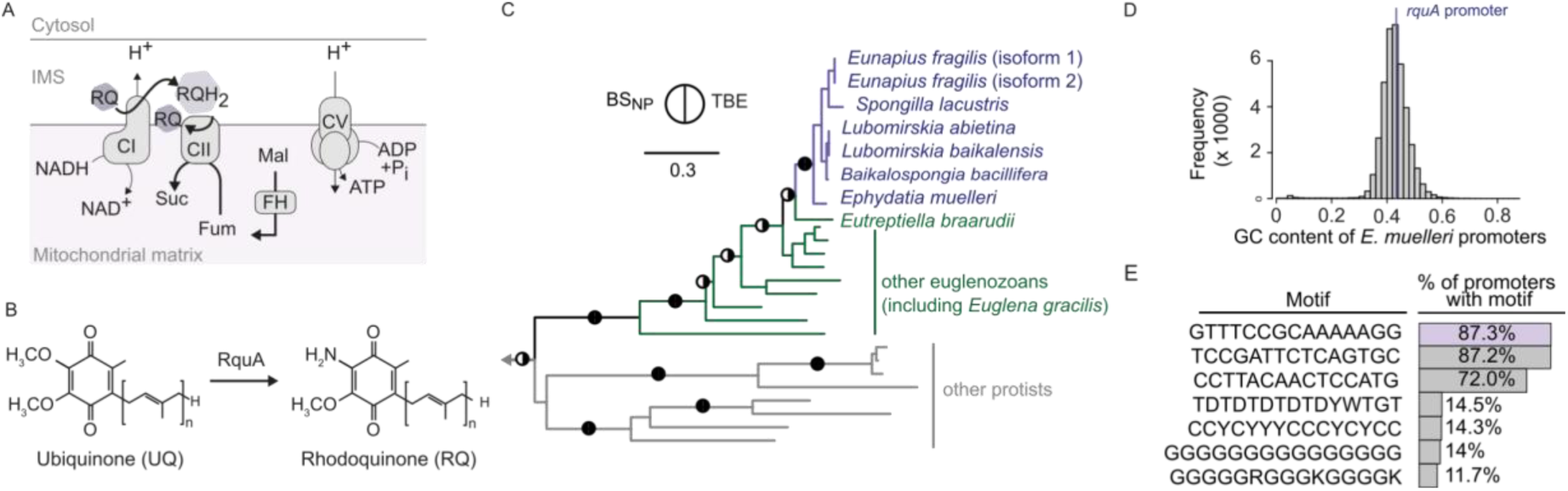
Metabolic function, lateral acquisition and genomic integration of *rquA* in freshwater sponges. **A.** Electrons derived from NADH are transferred to RQ via Complex I (CI), generating reduced rhodoquinol (RQH_2_). Malate is converted to fumarate (Fum) via fumarate hydratase (FH). RQH_2_ is subsequently oxidized by Complex II (CII) operating in reverse as a fumarate reductase, thereby reducing fumarate to succinate (Suc; modified from^3,13^). **B.** In some bacteria and protists, UQ is a precursor of RQ and this conversion is mediated by RquA (modified from^3,13^). **C.** Maximum-likelihood phylogeny of RquA proteins showing freshwater sponge homologs (purple) nested within a well-supported euglenozoa clade (green). Branch support values are indicated at nodes as nonparametric bootstrap (BS_NP_) and transfer bootstrap expectation (TBE). Filled black circles denote strongly supported nodes (BS_NP_ ≥ 95% and/or TBE ≥ 0.95). The scale bar corresponds to 0.3 substitutions per site. Non-euglenozoan protist sequences are shown in grey. **D,** Distribution of GC content across all annotated promoter regions in the *E. muelleri* genome (-1000 to +100 bp relative to the transcription start site). GC value of the *rquA* promoter (purple line). **E,** Genome-wide promoter motif frequencies (bars) and motif content of the *rquA* promoter (purple).

UQ biosynthesis is highly conserved across taxa and occurs in mitochondria via a multiprotein complex (Complex Q) that mediates substrate channeling through successive enzymatic reactions^15,16^. By contrast, distinct mechanisms for RQ biosynthesis have evolved across unicellular and multicellular organisms. In animals, RQ biosynthesis is initiated by a splice isoform of the prenyltransferase *coq2* (*coq2e*), which enables incorporation of a tryptophan-derived precursor (4-hydroxyanthranilate) into Complex Q, thereby diverting the UQ pathway towards RQ biosynthesis^14,17,18^. In contrast, bacteria and microbial eukaryotes generate RQ from UQ directly via the RquA protein, which facilitates the amination of the quinone ring^3,19^ (Fig. 1B).

Here, we reconstructed the phylogenetic distribution of RquA protein across prokaryotes and eukaryotes. Unexpectedly, we identified homologues of the microbial RquA within genomic and transcriptomic sequences of freshwater sponges (Demospongiae) and this correlates with RQ production in both wild and laboratory-grown sponges. Sponges (Porifera) represent one of the earliest-diverging metazoan lineages^20,21^ and inhabit environments characterized by fluctuating oxygen availability^22–25^. The presence of a microbial-type *rquA* gene in freshwater sponges provides evidence for the lateral acquisition of an anaerobiosis-associated metabolic gene in an animal lineage and reveals the contribution of LGT to metabolic adaptation in early-diverging metazoans.

### Freshwater sponges acquired RquA from protists via lateral gene transfer

To examine the distribution of RquA across prokaryotes and eukaryotes, we conducted sequence similarity searches using the *Pygsuia biforma* (Breviatea) RquA protein (GenBank: AKA62179.1) as a query against the EukProt database (v2). These searches unexpectedly recovered a homolog in the predicted proteome^26^ of the freshwater sponge *Ephydatia muelleri* (Demospongiae; *Em0002g590a*). Analysis of proximity ligation data showed continuous chromosomal interactions across the *Em0002g590a* locus with *cis* contacts extending into both flanking regions, consistent with chromosomal integration (Fig. S1). Chromatin profiling further revealed that the locus resides in a transcriptionally permissive chromatin environment, marked by promoter-associated H3K4me3 enrichment, characteristic of expressed endogenous genes (Fig. S1). Investigation of published transcriptomic datasets shows that *Em0002g590a* is consistently expressed across all developmental stages of *E. muelleri*^26^. Single-cell transcriptomic data^27^ further indicate that *rquA* expression is restricted to sclerocytes, archeocytes, and, our analysis shows also expression in choanocytes (Fig. S1) while it is undetectable in cystencytes, and basal pinacocytes^27^ (Transcript Per Million: TPM < 0.01; Table S1).

To further assess the distribution of RquA within Porifera, we performed additional homology searches against a dataset of sponge genomes and transcriptomes (Table S2). These analyses recovered RquA homologs exclusively from freshwater sponge lineages (demosponges), including *Lubromirskia abientina*, *Lubromirskia baikalensis, Baikalospongia bacillifera*, *Eunapius fragilis*, *Spongilla lacustris*, and *Ephydatia muelleri*, whereas no homologs were detected in marine conspecifics or in the transcriptome of the freshwater species *Ephydatia fluviatilis*. The absence of RquA in *E. fluviatilis* may reflect lineage-specific gene loss or incomplete sequencing of the transcriptome. All of the sponge RquA proteins possessed a predicted N-terminal mitochondrial targeting sequence, suggesting the proteins function in the mitochondrial compartment.

To evaluate the evolutionary origin of sponge RquA, we performed maximum-likelihood phylogenetic inference using RquA amino acid sequences retrieved from both eukaryotes and prokaryotes^3^. Phylogenetic reconstruction resolved the sponge RquA protein sequences as a well-supported clade that emerges from within a clade of sequences from euglenozoa protists. These closest relatives of the sponge RquA sequences derive from euglenozoa protists that inhabit marine/brackish (*Eutreptiella braarudii*)^28^ and freshwater habitats, including *Euglena gracilis*, a known RQ producer^29^ (Fig. 1C; Supplementary Data 1). This topology (Metazoa nested within Euglenozoa) is discordant with well-established phylogenetic relationships of the eukaryotic tree^30^, suggesting that *rquA* was acquired by an ancestor of freshwater demosponges from a euglenozoa-related protist via lateral gene transfer.

### Regulatory assimilation of *rquA* into the *Ephydatia muelleri* promoter landscape

To assess the sequence-level assimilation of the laterally acquired *rquA* gene within the *E. muelleri* genome, we compared the GC-content, k-mer frequency, and motif content of the *rquA* promoters to genome-wide promoter landscapes. The *rquA* promoter closely resembles endogenous promoters of core sponge genes, showing no deviation in GC content or k-mer composition from the genome-wide background (Fig. 1D; Fig. S2; Supplementary Data 2) and belonging to the predominant class of CpG-rich, TATA-less promoters. While most *E. muelleri* promoters possess up to seven sequence motifs, we could only detect the most common motif in the *rquA* promoter (Fig. 1E, purple). The promoter appears fully integrated into the host background at the level of base composition and CpG structure, while its atypical motif repertoire could indicate the gene’s more recent integration into the genome compared to resident genes.

### Yeast expressing sponge rquA synthesize RQ

To test whether the *E. muelleri* RquA (*Em*-RquA) is functional, we performed heterologous gene expression in *Saccharomyces cerevisiae* (that cannot naturally synthesize RQ) by placing the full-length coding sequence on an expression plasmid under the control of a galactose inducible promoter. Given that UQ and RQ biosynthesis occurs in the mitochondrion, we confirmed that *Em*-RquA, complete with its endogenous mitochondrial targeting sequence, is sufficient to direct the green fluorescent fusion protein *Em*-RquA-GFP to yeast mitochondria (Fig. S3; Supplementary Data 3).

We next assessed relative levels of quinones in yeast expressing the *Em*-RquA gene construct using quantitative mass spectrometry (Table S3). While uninduced and *Em*-RquA-expressing *S. cerevisiae* produced hexaprenyl species ubiquinone (UQ_6_)^31^, yeast cells expressing *Em-*RquA were found to contain RQ_6_ corresponding to 19.48 % of the total quinone pool. RQ_6_ was undetectable in uninduced controls (Welch’s *t*-test, *t* = -44.9, d.f. = 3.1, *p* < 0.001; Fig. 2A; Supplementary Data 4), indicating that expression of the sponge *rquA* gene is sufficient to confer RQ production in yeast.

**Fig. 2.**
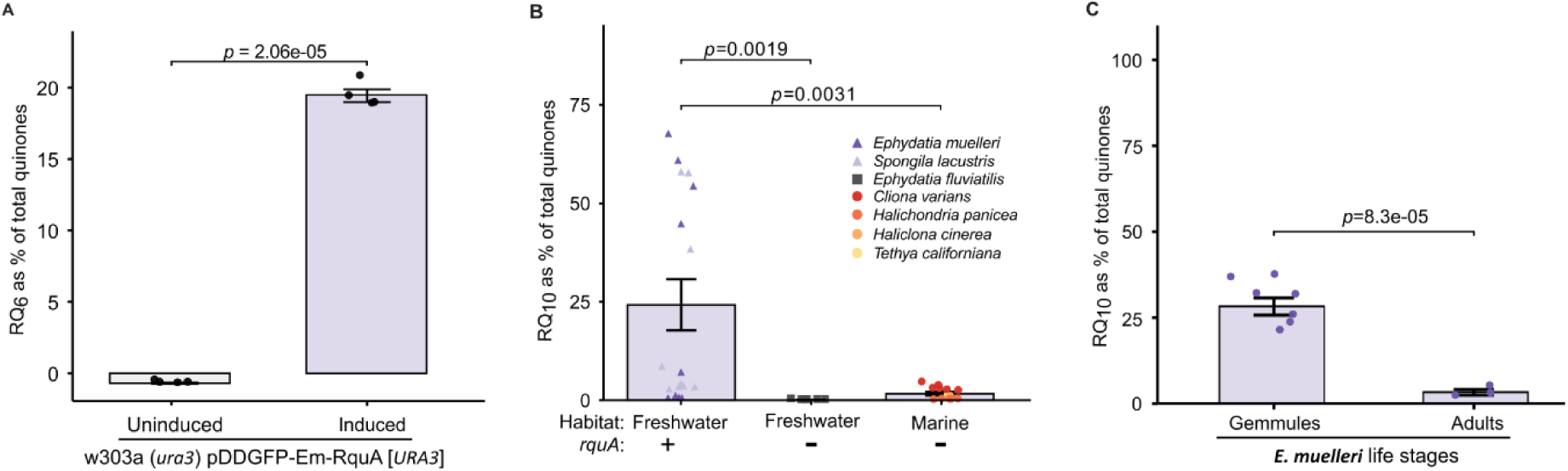
Detection of RQ in yeast expressing the *E. muelleri rquA* and in *rquA*-encoding freshwater sponges. **A.** Relative levels of RQ_6_ illustrated as a percentage of total quinones (UQ_6_ + UQ_6_H_2_ + RQ_6_) in yeast expressing *E. muelleri* RquA (*Em*-RquA) compared to uninduced controls. **B.** Relative levels of RQ_10_ illustrated as a percentage of total quinones (UQ_10_ + RQ_10_) across *rquA*-encoding freshwater sponges, freshwater sponges that do not encode *rquA*, and marine sponges. Bars represent mean ± s.e. Individual biological replicates per species are shown as circles with indicated colors, **C,** Relative levels of RQ_10_ detected illustrated as a percentage of total quinones (UQ_10_ + RQ_10_) in *E. muelleri* gemmules compared to adult tissue.

### RQ_10_ is the dominant rhodoquinone species in *rquA*-encoding freshwater sponges

To determine if *rquA*-encoding sponges contain RQ, we quantified UQ and RQ levels, in sponge specimens collected from freshwater and marine environments (Tables S4, S5). These taxa included two freshwater sponges encoding *rquA* (*E. muelleri* and *S. lacustris*), the freshwater sponge *E. fluviatilis*, which lacks detectable homologue for *rquA*, and four marine sponge species (*Halichondria panicea*, *Tethya californiana*, *Cliona varians* forma *varians,* and *Haliclona* cf. *xena*).

UQ_8_, UQ_9_, and UQ_10_ were detected in all sponges irrespective of habitat or *rquA* presence, with UQ_10_ being the most abundant (Fig. S4A-C). In contrast, RQ was consistently detected in freshwater sponges encoding *rquA,* with RQ_10_ and RQ_8_ representing the predominant RQ species (Fig. 2B; Fig. S4A-C). In *rquA*-encoding freshwater sponges, RQ_10_ accounted for 24.21% of the total quinone pool, a proportion significantly higher than that observed in *E. fluviatilis* that lacks *rquA* (Welch’s *t*-test, *t* = 3.72, d.f. = 16, *p* = 0.002) and in marine sponges (Welch’s *t*-test, *t* = 3.47, d.f. = 16.1, *p* = 0.003; Fig. 2B; Supplementary Data 5). Although most marine sponges lacked detectable RQ species, low levels of RQ_10_ were observed in *C. varians* forma *varians* (Fig 2B, Fig. S4C). Whether this RQ derives from the sponge or from its bacterial community cannot be determined with present data. In a laboratory setting, *E. muelleri* can be propagated from a dormant ‘gemmule’ stage 0 to an adult stage defined by the presence of a water pumping osculum. We isolated quinones from stage 0 gemmules and stage 5 adults and found a higher quantity of RQ_10_ in gemmules compared to adult tissue (Fig. 2C), suggesting stage-specific utilization of RQ in the animal (Welch’s *t*-test, *t* = 9.43, d.f. = 5.97, *p* < 0.001; Table S6).

### Hypoxia induces *rquA* expression without morphological alterations in the freshwater sponge *Ephydatia muelleri*

Central to sponge body organization are the choanocyte chambers, the osculum, and the aquiferous system, comprising incurrent and excurrent canals (Fig. 3A, 3B). To assess whether hypoxia induces morphological or cellular alterations, adult *E. muelleri* individuals reared under controlled laboratory conditions (∼20 °C, dark) were exposed to hypoxia (2% O_2_) and examined with microscopy. Light microscopy revealed no alterations in major structural features across all hypoxic time points examined (Fig. 3C; Supplementary Data 6). Consistent with this observation, confocal microscopy showed that choanocyte chambers retained normal morphology and organization following up to 72 h of hypoxic exposure (Fig. 3C; Supplementary Data 7).

**Fig. 3.**
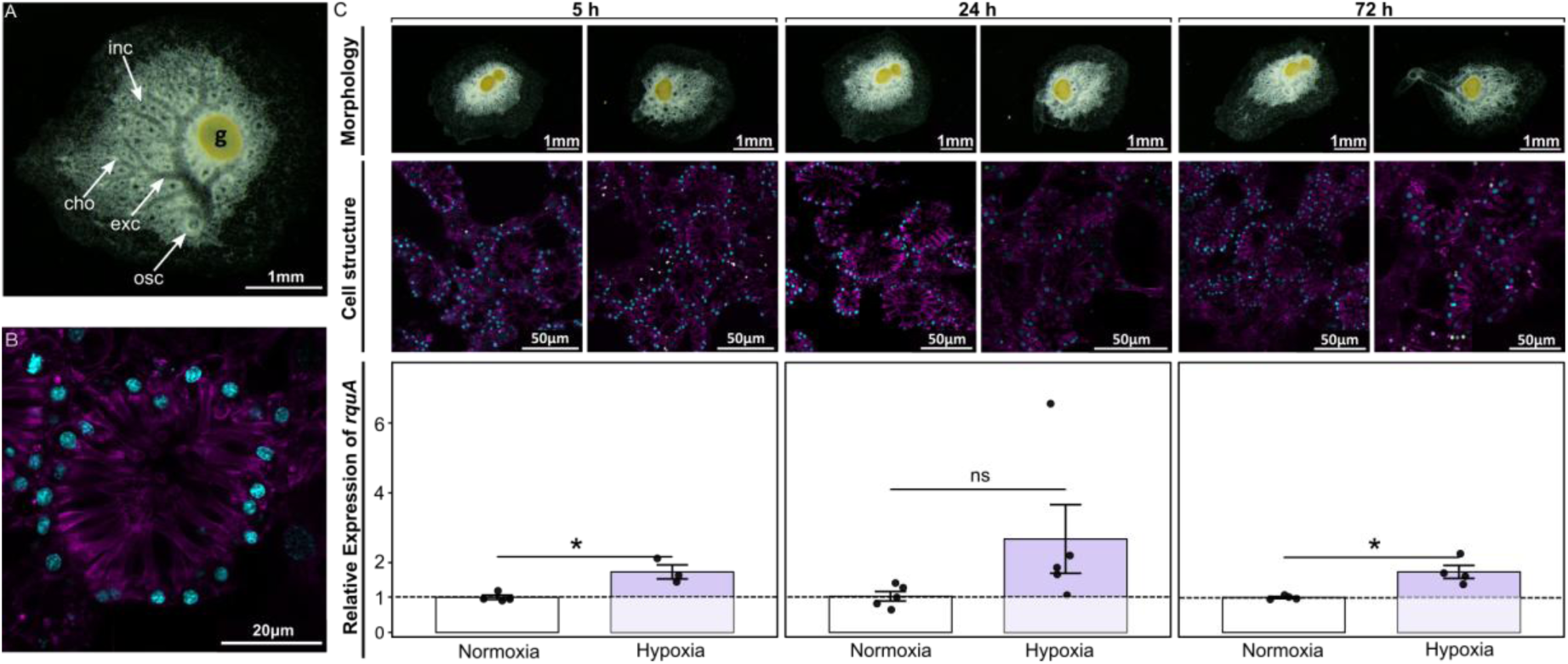
*E. muelleri* morphology and relative gene expression of *rquA* under normoxia (21% O_2_) and hypoxia (2% O_2_) after 5, 24, and 72 h. **A.** *E. muelleri* morphology annotated with the main sponge features, inc: Incurrent canal, exc: excurrent canal, cho: choanosome, osc: osculum, g: gemmule. **B.** *E. muelleri* cell-morphology of the choanocyte chambers. **C.** Whole-animal brightfield images (upper panel) showing overall body architecture and confocal images (middle panel) of choanocyte chambers (cell nuclei in cyan, actin in magenta, algae symbionts in green) reveal that chamber organization and morphology are preserved under hypoxia. Expression of *Em*-*rquA* (bottom panel) in *E. muelleri* was normalized to *ef1α* and *actin* and is presented relative to normoxic controls. Bars represent mean ± s.e. *P*-values were calculated using Welch’s *t*-test; *p<0.05.

To assess whether hypoxia elicits a transcriptional response in the absence of morphological change, we quantified *rquA* expression (Table S7). Expression levels were normalized to the reference genes *actin* and *ef1α* and calculated using the Pfaffl method^32^ which accounts for differences in primer efficiencies. The expression of *rquA* was significantly increased under hypoxia relative to normoxia (21% O₂) after 5 h (Welch’s *t*-test, *t* = -4.22, d.f. = 3.11, *p* = 0.034) and 72 h (Welch’s *t*-test, *t* = -5.00, d.f. = 3.46, *p* = 0.033), whereas the increase observed at 24 h was not significant (Welch’s *t*-test, *t* = -2.39, d.f. = 5.55, *p* = 0.058) (Fig. 3C; Supplementary Data 8).

### Freshwater sponges lack canonical UQ biosynthesis and convert exogenous UQ to RQ

Because UQ functions as the precursor of RQ biosynthesis in microbial systems^31,33^, we examined whether sponges encode the pathway required for UQ biosynthesis. In eukaryotes, UQ biosynthesis is mediated by a multi-protein pathway that includes enzymes catalyzing the core biochemical transformations of the isoprenoid tail and benzoquinone ring (Coq1-Coq7), as well as additional enzymes that contribute to the assembly, stability, and organization of the Q complex (Coq8 - Coq10)^15,16,34^.

To assess the distribution of these components in sponges, we surveyed homologs of Coq1-Coq10 across sponge genomes and transcriptomes, together with representative opisthokont model organisms (Table S8). Phylogenetic analyses using maximum likelihood of individual Coq protein sequences consistently placed sponge homologues among metazoan homologues, clearly separated from bacterial and plant homologues with strong support (UF ≥ 95; Supplementary Data 9). Across Porifera, we observed pronounced differences in the representation of UQ biosynthetic genes. Freshwater sponges consistently lacked homologues of Coq1, Coq2, Coq4, Coq5, Coq6. In contrast, Coq8-10, and the terminal hydroxylase Coq7, were recovered across all examined freshwater taxa (Fig. 4A; Table S8). Conversely, marine sponges exhibited greater retention of genes involved in UQ biosynthesis. Members of the classes Calcarea and Homoscleromorpha possessed homologs of all annotated Coq enzymes, as did marine demosponges of the subclasses Keratosa and Verongimorpha. Within Heteroscleromorpha, however, Coq enzyme representation was more heterogeneous, with several orders lacking subsets of core catalytic components despite retaining genes associated with assembly functions.

**Fig 4.**
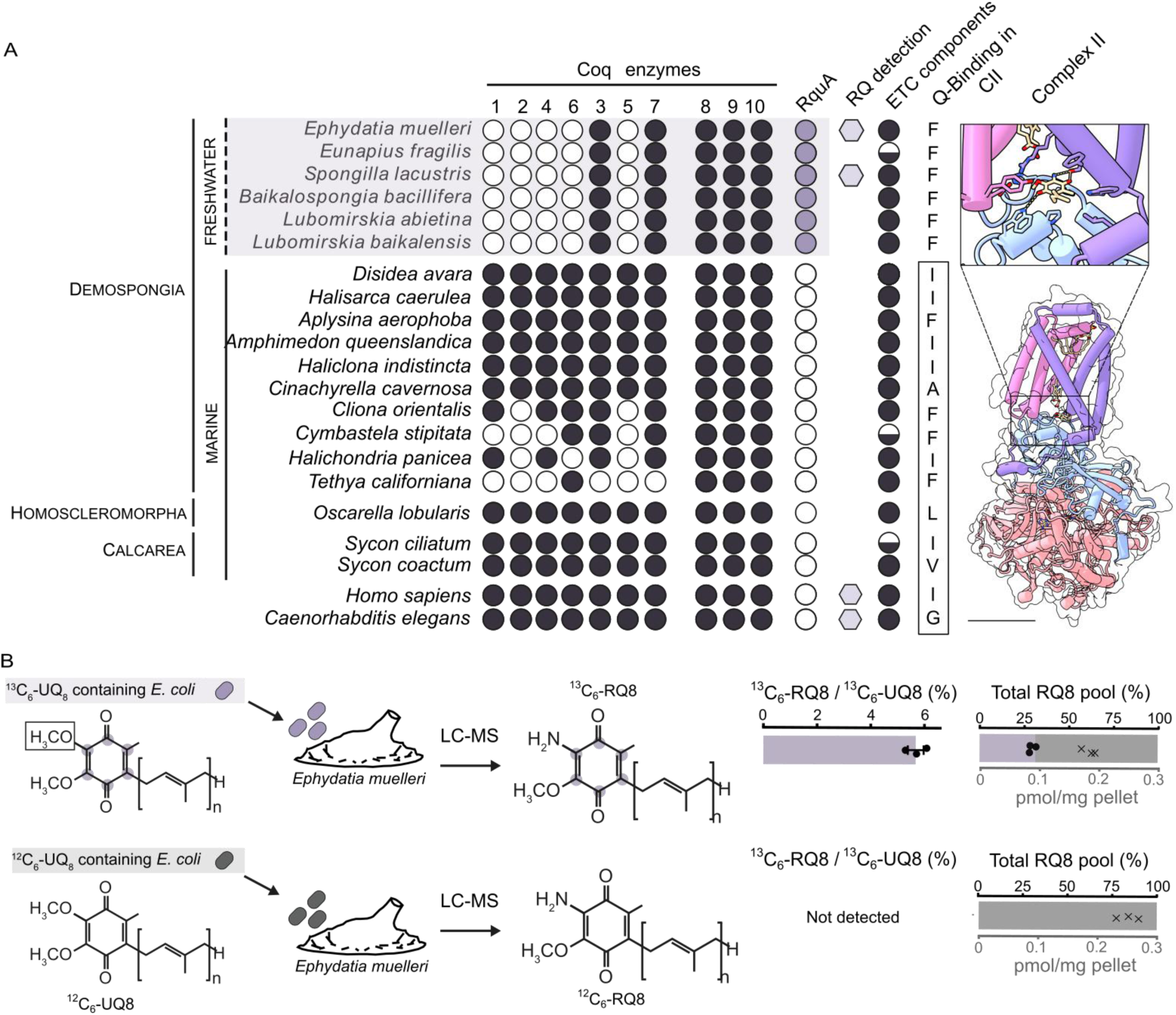
**A,** Presence (black circles) and absence (white circles) of Coq enzyme homologues and RquA (purple circles) across sponge lineages and representative opisthokonts, Complex Q components are ordered in their proposed reaction sequence (Coq1-10). Hexagons denote taxa in which RQ has been experimentally detected. Half circles for the ETC components indicate that not all subunits were detected; All ETC components are presented in Fig. S5 in details. Complex II binding site indicates residue variation in site 5 (quinone-binding site in Complex II). **B,** Conversion of dietary ^13^C_6_-UQ_8_ to ^13^C_6_-RQ_8_ expressed as the ratio ^13^C_6_-RQ_8_/^13^C_6_-UQ_8_ across treatments, and RQ_8_ isotopologues as a percentage of the total RQ_8_ pool. Data points (circles and crosses) indicate absolute RQ_8_ abundance (pmol/mg pellet) of ^13^C_6_-RQ_8_ and ^12^C_6_-RQ_8_, respectively. Purple: ^13^C_6_-RQ_8_ labeled *E. coli*; grey: unlabelled *E. coli*.

The absence of several core catalytic components of the canonical UQ biosynthetic pathway in freshwater sponges suggests that these taxa may rely on exogenous resources of UQ or UQ biosynthesis intermediates for RQ biosynthesis. To determine whether freshwater sponges can utilize exogenous UQ as a substrate to synthesize RQ, we fed laboratory-reared *E. muelleri* adults with ^13^C_6_-UQ_8_-labelled *Escherichia coli*, and quantified quinone species by LC–MS^33^. We detected ^13^C_6_-RQ_8_ only in sponges fed with ^13^C_6_-UQ_8_-labelled *E. coli*, whereas no labelled RQ_8_ was observed in sponges fed unlabelled *E. coli* (Fig. 4B, Table S9; Supplementary Data S10). Although the abundance of ^13^C_6_-RQ_8_ was low, its restricted occurrence in the labelled treatment supports that dietary UQ_8_ can be converted into RQ_8_ *in vivo* in freshwater sponges.

### RQ biosynthesis and utilization is associated with few molecular changes to respiratory chain components

RQ biosynthesis in eukaryotes often coincides with reduction of ETC components^13^ or expression of RQ-specific subunits of CI and CII^18^. To determine if similar gene gain or loss events have occurred in *rquA*-encoding sponges compared to marine sponges, we performed a comparative genomic investigation of respiratory components across representative Porifera. We found the core components of CI-CV are conserved across marine and freshwater sponges (Fig. S5), suggesting all have canonical respiratory capabilities. The missing subunits from *Cymbastela stipitata, Sycon ciliatum,* and *E. fluviatilis* are attributed to incomplete datasets and are likely spurious absences. Unlike other RQ-utilizing animals^13^, all quinone interacting subunits were present as single copies, suggesting that the respiratory complexes of worms and sponges have evolved independent strategies to utilize RQ.

Given the central role of CII in quinone-dependent electron transfer^13^, we examined conservation of amino acid substitutions within the quinone-binding region of this complex^35–39^ across metazoan available on the interpro database. Most metazoan that are predicted to use UQ as their primary quinone, possess a non-polar amino acid (I/A/V/L) in the quinone binding pocket, including most marine sponges (Fig. 4; Fig. S6). However, metazoan and other eukaryotes known or predicted to produce RQ have phenylalanine (e.g., freshwater sponges, platyhelminths, *Quaeritorhiza haematococci* [fungi], and *Paramoeba* species [amoebozoa]) or glycine (e.g., nematodes) in the homologous site, hinting that RQ utilization might correlate with changes in substrate binding site evolution. Phenylalanine is also present in the SdhC proteins of non-*rquA*-encoding marine sponges *Aplysina aerophoba*, *Cliona orientalis*, *Cliona varians* and *Cymbastella stipitata,* suggesting there might be cryptic RQ production in these organisms. Although phenylalanine introduces an aromatic side chain relative to the aliphatic residues, all substitutions at this site are hydrophobic, and phenylalanine is not restricted to known *rquA*-encoding lineages or RQ utilizers. The uniform presence of phenylalanine in freshwater sponge taxa may reflect lineage-specific conservation. It is unknown what the biological implication of these substitutions are, however, because quinone rings are capable of non-covalent interactions with aromatic side chains (i.e., π-stacking), it is possible this phenylalanine participates in π-mediated intermolecular interactions^40^.

To assess whether variation at this SdhC residue affects quinone binding, we performed *in silico* docking predictions and estimated binding affinities of UQ and RQ species in Complex II (CII) from *E. muelleri* and *Homo sapiens*. There is no significant difference in predicted binding affinity between UQ and RQ for *E. muelleri* and *H. sapiens* (Fig. S5; Supplementary Data 12). This suggests that variation at this SdhC residue alone is insufficient to confer quinone-binding specificity. In *E. muelleri,* RQ_1_ shows higher binding affinity than RQ1H2, indicating that the oxidized form of quinone might be better accommodated into the binding pocket. It should be noted that local sidechain conformation may have an accuracy issue in these predictions, which has been reported in some protein language models^41^.

## Discussion

The presence of a microbial-type RquA in freshwater demosponges reveals a previously unrecognized route by which animals can acquire anaerobic metabolic capacity. Rather than evolving RQ biosynthesis through modification of the UQ biosynthesis pathway like in other metazoans^14,18^, freshwater sponges appear to have incorporated a microbial gene via LGT. The restricted phylogenetic distribution of *rquA* in freshwater sponges is consistent with a single acquisition event. This event likely occurred after the transition of demosponges into freshwater habitats (∼250–300 Mya) and before the Cenozoic diversification of extant freshwater taxa (∼15–30 Ma)^26^. Given that freshwater habitats are characterized by fluctuating oxygen availability and recurrent hypoxia^42^, it is likely that oxygen limitation may have been a persistent selective pressure during this period, favoring the retention of anaerobic metabolic traits. More broadly, these findings indicate that RQ biosynthesis in animals does not follow a single evolutionary trajectory. The detection of RQ in taxa lacking both known strategies for RQ production (*rquA* and *coq2* splice variants), including the marine sponge *C. varians* (this work) and vertebrate tissues^43^, suggests that multiple biosynthetic or acquisition pathways may exist. Together, these observations support a model in which anaerobic quinone metabolism in animals has evolved through diverse mechanisms, including lateral gene transfer and endogenous pathway modification.

Because UQ serves as the precursor for RQ in the microbial pathway^3,19^, animals encoding *rquA* would be expected to retain the full ubiquinone biosynthetic pathway. However, freshwater sponges show a reduced representation of core UQ biosynthetic genes, indicating that *de novo* UQ synthesis is not universal across animal lineages. Instead, RQ production in these taxa appears to rely on exogenous quinones, consistent with their filter-feeding ecology and the high organic content of many freshwater systems. Some heterotrophic protists that possess RQ and *rquA* completely lack a UQ biosynthesis pathway^3^, suggesting that dietary UQ is sufficient to allow for maintenance of RQ biosynthesis machinery. UQ biosynthetic genes seem to be inconsistently represented across marine sponges, suggesting that neither habitat nor the presence of *rquA* alone predicts retention of the canonical pathway. Together, these observations indicate that quinone metabolism in animals is more variable than previously assumed.

The cellular organization of sponges provides a permissive context for laterally transferred genes and their long-term retention. Their feeding mode, in which choanocytes phagocytose bacteria and picoplankton, exposes internal cell types to microbial DNA^44–46^. In addition, the absence of strict germline segregation increases the likelihood that acquired genes become heritable. In freshwater sponges, *rquA* is predominantly expressed in archeocytes, mesohyl-resident cells that contribute to both somatic tissues and gametogenesis^27,44,47^. Because these cells function as pluripotent types linking somatic maintenance and reproduction, the expression of *rquA* in these cells provides a direct route for transmission of the acquired gene across generations. This pattern suggests that long-term retention of laterally transferred genes in early animals may depend less on the initial transfer event than on their subsequent integration into cell lineages with developmental and reproductive continuity.

Morphological and cellular evidence indicates that freshwater sponges survive in hypoxic conditions. However, this tolerance is not unique to *rquA*-encoding taxa, as many marine sponges lacking the gene also survive prolonged hypoxia^22–25^. Such marine taxa also experience transient internal hypoxia during periodic contractions^48^, arguing against a primary role or RquA in contraction-related physiology. Instead, the retention of *rquA* is more plausibly linked to specific life-history stages, such as gemmules, that endure prolonged anoxic conditions^49^. Consistent with this interpretation, RQ levels were elevated in gemmules relative to adult tissues, likely reflecting the higher abundance of archeocyte-like cells. This pattern is further supported by the cellular expression profile of *rquA*, which is predominantly detected in archeocytes and sclerocytes, the two principal cell types that compose gemmules^50^. Together, these observations suggest a link between *rquA* expression, RQ accumulation, and the cellular lineages responsible for gemmule formation and persistence. More broadly, our findings indicate that anaerobic metabolic innovation in animals can arise through the acquisition of microbial genes expressed in specific cell types, thereby linking LGT to life-history adaptations. In freshwater sponges, this integration enables the utilization of an anaerobic quinone system that likely facilitates survival under hypoxic conditions, showing how horizontal gene acquisition can reshape metabolic evolution in early animals.

## Methods

### Phylogenetic reconstruction of bacterial and eukaryotic RquA homologs

To identify RquA homologs we queried the *Pygsuia biforma* RquA protein sequence (GenBank: AKA62179.1) against the EukProt database^51^ using blastp with an e-value cut-off of 1e-10. This search yielded an RquA homolog in the conceptual proteome of the freshwater sponge *Ephydatia muelleri* (BioProject: PRJNA579531^26^). This protein sequence was subsequently used as a query in tblastn homolog searches (e-value threshold: 1e-5) against a sponge dataset with genomic and transcriptomic resources (Table S2). Among opisthokonts, RquA was detected in the fungus *Quaeritorhiza haematococci* (JADGJF010000989.1, scaffold_989).

Identified RquA homologs from freshwater demosponges (*Eunapius fragilis, Spongilla lacustris, Baikalospongia bacillifera, Lubomirskia baikalensis,* and *Lubomirskia abietina*) were added to previously published RquA dataset^3^, which was further supplemented with homologs from Euglenozoa [per. communication Gordon Lax (U. British Columbia) (*Dryad Data*^52^). Amino acid sequences were aligned using MAFFT v7.490^53^ and ambiguously aligned residues were removed using BMGE 209 v1.12^54^.

Maximum-likelihood phylogenetic inference was performed with IQ-TREE v2.2.2.6^55,56^. The initial phylogeny was inferred under the WAG+C40+F+G mode of evolution (selected using model selection: -mrate G -mset LG, LG+C10, LG+C20, LG+C30, LG+C40, JTT, JTT+C10, JTT+C20, JTT+C30, JTT+C40, WAG, WAG+C10, WAG+C20, WAG+C30, WAG+C40, LG4X, LG4) with 1000 ultrafast bootstraps and 1000 SH-aLRT replicates. The final tree was inferred using the posterior mean site frequency (PMSF^57^) approximation using the ultrafast bootstrap tree as a guide tree to calculate the site specific frequencies and 100 non-parametric bootstraps (-b 100), and the transfer bootstrap expectation (-tbe)^58,59^ were inferred.

### Hi-C evidence for chromosomal embedding of Em0002g590a in the *Ephydatia muelleri* genome

To assess the integration of *Em0002g590a* in the chromatin contact landscape, we used previously published Micro-C proximity ligation data generated from sorted *E. muelleri* choanocytes^60^. Raw Micro-C data were mapped to the reference genome assemblies GCA_013339895.1^26^ and GCA_049114765.1^60^ as described here https://github.com/sebepedroslab/early-metazoa-3D-chromatin. Briefly, raw reads were aligned to the reference genome using bwa-mem (v.0.7.17-r1188) with the option -SP5M. Mapped reads were sorted, paired and deduplicated using pairtools (v.0.3.0)^61^. We also discarded reads mapping within 200 bp to adjacent nucleosomes. Contact matrices were generated and normalized using cooler (v.0.8.11)^62^, and chromatin interaction profiles were visualized with Coolbox (v.0.3.8)^63^. Chromatin profiling ChIP-seq data and expression 3’ RNA-seq (MARS-seq) data were generated previously^60^. Raw ChIP-seq data was mapped to reference genomes using bwa mem (version 0.7.17-r1188), and 3’ RNA-seq raw reads were aligned to the reference genomes using STAR (v.020201)^64^. Genome coverage profiles were generated with deeptools (v. 3.1.3)^65^.

### Promoter architecture and regulatory context of *rquA*

We analyzed the genomic regulatory context of *rquA* using the *Ephydatia muelleri* genome assembly and structural annotation obtained from EphyBase (https://spaces.facsci.ualberta.ca/ephybase/)^26^. Transcription start sites (TSSs) were extracted from the GFF3 annotation as strand-specific 5′ ends of annotated transcripts. Promoter regions were extracted as −1000 bp upstream to +100 bp downstream of each TSS in a strand aware manner using bedtools (v2.31.1)^66^.

We computed GC content for all promoter sequences using bedtools nuc and summarized in R (v4.1.2). The GC-content of the *rquA* promoter was compared to the genome-wide promoter distribution to assess compositional similarity. To further characterize promoter sequence composition, all possible 4-mers over {A, C, G, T} were counted and converted to relative frequencies, yielding an N × 256 promoter-by-k-mer matrix. K-mer profiles were constructed using custom Python scripts (Python 3.9.16, Biopython 1.81, pandas 2.1.3, numpy 1.26).

Principal component analysis (PCA) was performed using scikit-learn (v1.6.1) on centered 4-mer frequency vectors to visualize large-scale compositional patterns and to project the *rquA* promoter into the same space. Because k-mer frequencies are compositional data constrained to sum to one, PCA was also performed after a centered log-ratio (CLR) transformation using a pseudocount (ε = 10^-8^) to avoid undefined logarithms. The CLR transformation corrects for the constant-sum constraint and allows the analysis to reflect relative differences among k-mers. In both cases, the *rquA* promoter was projected into the same compositional space for comparison.

We conducted *de novo* motif discovery using MEME v5.5.9^67^ (ZOOPS model; DNA mode; 10 motifs; motif widths 6–15 bp; reverse complement enabled). Genome-wide motif occurrences were identified using FIMO v5.5.9^68^ (*p* ≤ 1×10^-3^) and output was summarized to a promoter x motif count matrix. Further, motifs were annotated by comparison to reference databases using TOMTOM v5.5.9^69^ (*q* ≤ 0.05). Custom Python scripts were used to quantify (i) genome-wide motif frequencies, (ii) motif presence/absence in the *rquA* promoter, (iii) depletion of common motifs, and (iv) total motif counts relative to background, including z-score analysis.

To assess core promoter features, we screened all promoters for TATA-box motifs using the JASPAR TBP PWM (MA0108.1)^70^ as FIMO input (*p* ≤ 1×10^-4^). Promoters containing at least one significant match were classified as TATA-positive. The *rquA* promoter was evaluated using the same criteria.

Finally, we analyzed CpG content by computing observed-to-expected CpG ratios for all promoters.

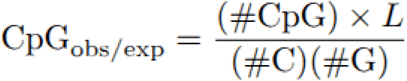

We further identified CpG islands using the Gardiner–Garden and Frommer criteria^71^ (GC ≥ 50%, CpG O/E ≥ 0.6, length ≥ 200 bp). Sliding-window CpG density profiles (200 bp window, 20 bp step) were computed genome-wide, and corresponding metrics were extracted for the *rquA* promoter using custom Python scripts.

### Heterologous gene expression and fluorescent microscopy of *Saccharomyces cerevisiae*

The full-length *E. muelleri rquA* coding sequence (*Em-rquA; Em0002g590a*) was codon optimized for expression in yeast and synthesized in the pUC57 plasmid (Genewiz). The optimized sequence was cloned into the pDDGFP_LEUD expression vector (Addgene #58352), under the control of the galactose-inducible GAL1 promoter. The resulting construct was transformed into *Saccharomyces cerevisiae* W303-1A strain^72^ using the S.c. EasyComp Transformation kit (Invitrogen). Transformants were selected on synthetic defined (SD) plates lacking ura and verified by colony PCR.

For fluorescence microscopy, transformants were grown overnight at 30 °C in SD-ura medium supplemented with 2% glucose to repress GAL1-driven expression. Cells were harvested by centrifugation (1500 x g for 5 min), washed in PBS, resuspended in pre-warmed SD-ura medium containing either glucose or galactose, and incubated at 30 °C for 4 – 6 hours to repress or induce *Em*-RquA–GFP expression, respectively. Wild-type W303-1A cells (grown in SD+ura medium) and cells carrying the empty pDDGFP_LEUD plasmid were included as controls.

Mitochondria and nuclei were stained with Mitotracker Red CMXRos (0.2 nM, 20 min) and DAPI (11 nM, 10 min), respectively. Cells were washed in PBS, mounted on 1% agarose pads, and imaged using a Zeiss Axio Imager Z2 microscope equipped with a Plan-Apochromat 100×/1.40 Oil Ph3 M27 objective. Images were acquired using excitation/emission wavelengths of 587/610 nm (MitoTracker), 488/509 nm (GFP), and 353/465 nm (DAPI). Exposure times were 100 ms (MitoTracker), 500 ms (GFP), and 170 ms (DAPI). Images were processed using linear adjustments (e.g., brightness/contrast) and deconvolution in Zeiss Zen software. Further processing was performed using ImageJ.

To prepare cell biomass for quinone analysis, overnight yeast cultures (W303-1A(ura) pDDGFP_LEUD-EmRquA) that were propagated in SD-ura+glucose were used to seed fresh cultures into SD-ura+glucose (uninduced) or SD-ura+galactose (to induce *Em*-RquA expression) for 8 h at 30 °C (200 rpm; 8 h). After 8h of incubation, cells were harvested by centrifugation (1,500 × g, 10 min), and stored at −80 °C until quinone extraction.

### Environmental sampling of natural marine and freshwater sponges

To characterize quinone composition across sponge taxa, we collected freshwater and marine sponges from multiple geographic locations (Tables S4, S5). Marine species included *Cliona varians forma varians* (n = 6), *Halichondria panicea* (n = 5), *Tethya californiana* (n = 1), and *Haliclona cf* xena (n = 1). Freshwater taxa included the *rquA*-encoding species *Ephydatia muelleri* (Canada, n = 4; Sweden, n = 5) and *Spongilla lacustris* (Canada, n = 3; Sweden, n = 5), as well as the non–*rquA*-encoding species *Ephydatia fluviatilis* (Sweden, n = 6). Specimens collected in Canada were identified morphologically by S. Leys (University of Alberta), whereas Swedish specimens were identified by sequencing the *cytochrome oxidase I* (*COI*) mtDNA marker^73^. Samples of *C. varians* forma *varians* were obtained from material collected in the Caribbean Sea in 2014^74^, flash frozen in liquid nitrogen and subsequently stored at −80°C.

To compare quinone composition across developmental stages, quinones were quantified from frozen gemmules (stage 0) and post-hatching adult individuals (stage 5) of *E. muelleri*. Gemmules were induced to hatch and reared *in vitro* in Petri dishes in the dark at ∼20°C. Sponges were sampled at developmental stage 5 (approximately 6 d post hatching), corresponding to fully developed individuals with a functional aquiferous system, differentiated choanocyte chambers, and a fully formed osculum (Fig. 3A). Sponge tissue was dislodged from the Petri dish using gentle scraping, transferred to a microcentrifuge tube, and collected by centrifugation in a low-speed centrifuge (1-2 s), and stored at −80 °C until quinone extraction.

### Quinone extraction and liquid chromatography-mass spectrometry (LC-MS) analysis

Quinone extraction and LC–MS analysis were performed using the same protocol for all samples. Quinones were extracted using a two-phase liquid–liquid procedure. Briefly, pellets were resuspended in ultrapure water (0.5 mL), supplemented with zirconium oxide beads for mechanical disruption, and complemented with UQ_3_ (30 pmol) as an internal standard. Quinone solubilization was facilitated by ethanol-assisted extraction at 70 °C, followed by partitioning into hexane. Organic phases were pooled, dried under nitrogen gas (Organomation, Berlin, MA), and resuspended in LC–MS–grade acetonitrile (100 µL). Extracts were analyzed immediately to minimize oxidative degradation.

UQ and RQ species with six, eight, nine and ten isoprenoid tail lengths were quantified by LC–MS using a triple quadrupole mass spectrometer (Waters Xevo Cronos TQ) coupled to reversed-phase chromatography on a pentafluorophenyl column (Phenomenex Luna PFP[2], 50 × 2 mm, 3 µm, 100 Å). Separation employed a binary solvent system of water and acetonitrile, each containing 0.1% formic acid, with the following linear gradient: 0–3.5 min (30:70), 3.5–3.75 min (30:70 to 2:98), 3.75–7.25 min (2:98), 7.25–7.5 min (2:98 to 30:70), and 7.5–9 min (30:70). Mass spectrometric detection was carried out in positive electrospray ionization mode (capillary voltage 0.6 kV, desolvation temperature 600 °C, desolvation gas flow 1,000 L h⁻¹). Quantification was achieved using multiple reaction monitoring (MRM), with compound-specific precursor–product ion transitions and optimized cone and collision voltages listed in Table S10. Yeast samples were analyzed using transitions targeting UQ_6_, UQ_6_H₂, and RQ_6_, whereas sponge samples were analyzed using transitions specific to UQ_8–10_ and RQ_8–10_. UQ_3_ (internal control) was monitored in all cases. Samples were maintained at 12 °C prior to injection and chromatographic separations were conducted at 25 °C.

### Quinone quantification and data analysis

Data was further processed using Waters MassLynx and TargetLynx software (v. 4.2) for peak detection, integration, and quantification. Quantification of UQ and RQ species was performed using external calibration curves generated from quinone standards (UQ_10_, RQ_10_, UQ_9_, RQ_9_, UQ_6_, and UQ_3_). Where RQ standards were unavailable, quantification was based on the corresponding UQ calibration curve using empirically determined response correction factors. Peak areas were normalized to the internal standard UQ_3_ and converted to molar quantities (pmol), which were further normalized to pellet biomass (pmol/mg pellet). In yeast, UQ_6_, UQ_6_H₂, and RQ_6_ were quantified. Because RQ_6_ exhibited a higher mass spectrometric response relative to UQ_6_, RQ_6_ values were corrected using a response factor of 2.3 prior to downstream analysis (Table S3). Negative RQ_6_ values in control samples arose as artifacts of dilution and calibration curve limitations and reflect trace levels (<1 pmol RQ_6_/mg pellet). Total UQ abundance in yeast was calculated as the sum of oxidized (UQ_6_) and reduced (UQ_6_H_2_) form. In sponge samples, UQ_8-10_ and RQ_8-10_ were quantified.

For all samples, quinone abundances were assessed for normality using the Shapiro–Wilk test and for homogeneity of variance using Levene’s test. Differences between groups were evaluated using Welch’s *t*-tests, comparing (i) wild-type yeast and yeast expressing *E. muelleri* RquA, (ii) *rquA*-encoding versus non-*rquA*-encoding freshwater sponges, (iii) *rquA*-encoding freshwater sponges versus marine sponges, and (iv) gemmules versus adult sponge tissue of *E. muelleri*. To assess the contribution of RQ to the total quinone pool, RQ representation was calculated as RQ / (UQ + RQ) for each quinone species and differences among the groups were assessed with Welch’s *t*-tests. All statistical analyses were performed in R (v4.2.2).

### Assessment of phenotypic responses to hypoxia in *Ephydatia muelleri*

Frozen gemmules of the freshwater sponge *E. muelleri* (O’Connor Lake, Canada) were thawed at room temperature, washed 5 times in dH_2_O to remove residual of DMSO and placed at 4 °C in the dark for priming. After 1 week, gemmules were transferred to Petri dishes containing 1X Strekal’s medium (Cation concentration: MgSO_4_·7H_2_O, 2 mg L⁻^1^; CaCO_3_, 2 mg L⁻^1^; Na_2_SiO_3_·9H_2_O, 0.5 mg L⁻^1^; KCl, 0.5 mg L⁻^1^). Cultures were maintained at 20 °C in the dark to induce hatching. At 6 d post-hatching, stage-5 adult sponges were transferred to a controlled hypoxia chamber, where oxygen concentration was set to 2% O_2_ (v/v). Oxygen levels were regulated using an oxygen regulator (ProOX P110, BioSpherix) coupled to N_2_ gas. Control sponges were maintained under normoxic conditions (21% O_2_, v/v). Sponges were imaged at 5 h, 24 h, and 72 h following hypoxia exposure using a stereomicroscope (Olympus SMZ1) equipped with a digital camera (E3CMOS). Overall body architecture, canal organization, and osculum structure were assessed qualitatively.

For cellular-level morphological analysis, sponges from the same batch were plated in 35 mm glass bottom dishes (#81158; Micromedic), and 6-d post hatching were exposed to either normoxia (21% O_2_) or hypoxia (2% O_2_) for 5 h, 24 h, and 72 h. After the exposure time, sponge tissues were fixed in 4% paraformaldehyde and incubated overnight at 4 °C. Tissue was washed twice in 1x PBS, permeabilized with 0.1% Tween20 in 1x PBS for 1 h, and incubated in the dark for 45 min in 0.5 ml PBS containing DAPI (final concentration: 1 μg/mL, Sigma Aldrich) and Atto 565 Phalloidin (final concentration: 24 μM, Sigma-Aldrich). Finally, sponges were washed 3 times in PBS and imaged using a Leica SP8 DLS confocal microscope using a 63x/1.3 Glycerol Objective. The LAS X and Fiji^75^ software was used for image analysis and visualization.

### Quantification of *rquA* gene expression under hypoxia in *Ephydatia muelleri*

Stage-5 sponges were exposed to either hypoxia (2% O₂, v/v) or normoxia (21% O₂, v/v) for 5 h, 24 h, or 72 h as described above. Sponge tissue was harvested by gently scraping sponges from the culture surface using a sterile pipette tip, transferred to RNAlater (Thermo Fisher Scientific), and stored at −80 °C until processing. At least four biological replicates per treatment and time point were collected, each consisting of pooled tissue from ∼15-20 sponges. RNAlater was removed by collecting the sponge tissue by a brief centrifugation and removing residual RNAlater by pipetting. Total RNA was isolated using TRIzol^TM^ (Thermo Fisher Scientific) followed by column purification using the Animal Tissue RNA Purification Kit (Norgen Biotek, Canada), with a final elution volume of 35 µL. Residual genomic DNA was removed by DNase I treatment (Thermo Fisher Scientific). RNA integrity was assessed qualitatively by electrophoresis on a 1% agarose gel, where intact 18S and 28S rRNA bands were observed, and RNA concentration was quantified using the Qubit RNA HS Assay (Thermo Fisher Scientific). An amount of 300 ng of total RNA was reverse-transcribed into cDNA using the iScript cDNA Synthesis Kit (Bio-Rad) with random primers.

Quantitative PCR (qPCR) was performed using the SsoAdvanced Universal SYBR Green Supermix (Bio-Rad) with *E. muelleri rquA*–specific primers (2591F: 5′-TGTCGTGTGGGTGGAGTA-3′; 2887R: 5′-CTCAATTTCTGTATGCCACATCC-3′). qPCR cycling conditions consisted of an initial denaturation at 95 °C for 3 min, followed by 40 cycles of 95 °C for 15 s and 56 °C for 15 s, with a melt-curve analysis performed at the end of each run to confirm amplification specificity. Expression levels were normalized to the housekeeping genes *ef1α* (F: 5′-GCGGAGGTATCGACAAGCGT-3′; R: 5′-AGCGCAATCGGCCTGTGAG-3′) and *actin* (F: 5′-CGTTCTTCCCCACGCCATC-3′; R: 5′-TCGCTCGGCAGTGGTGGT-3′). Relative expression was calculated using the Pfaffl method^32^. Primers were designed using Primer-BLAST software^76^. The primer efficiency was determined using standard curve method^77^. All qPCR reactions were performed in three technical replicates using a Bio-Rad CFX96 Real-Time PCR Detection System.

### Identification of Coq homologues across Metazoan

#### Reference queries and initial homology searches

Homologues of Coq1-10 were initially identified using profile-based searches. Hidden Markov model (HMM) profiles for each Coq family were downloaded from the NCBI Conserved Domain database and used as queries in *hmmsearch* (HMMER v3.1b2) against a custom database comprised of sponge genomes/transcriptomes and genomes of model organisms (Table S8). Sequences retrieved from the HMM searches were subsequently subjected to sequence similarity-based searches. Each candidate was used as a query in tblastn searches against the *nr* database with an e-value cutoff 1 x 10^-5^, and the top 100 hits were retained. Sponge sequences for which the majority of top hits (>80%) corresponded to bacterial homologs were considered likely contaminants and removed from further analyses.

To place sponge Coq candidate proteins in a broader evolutionary context, prokaryotic homologues were also collected. While some prokaryotic sequences were recovered directly through searches with Ubi-family HMM profiles, most were obtained via blastp searches against *nr* using the eukaryotic sequences retrieved by the HMM searches restricting to prokaryotes. Prokaryotic homologues were retained for downstream phylogenetic analyses to facilitate the detection of potential contamination in sponge resources. Eukaryotic and prokaryotic homologues were combined for phylogenetic reconstruction. For each Coq family (Coq1–Coq10), a comprehensive sequence set was assembled consisting of (i) sequences from the selected sponge species, (ii) sequences from the selected model organisms, and (iii) the top 100 *nr* hits retrieved for each of these queries. Because these searches produced highly redundant datasets with substantial overlap among hit lists, duplicate and near-identical sequences were removed prior to alignment. The resulting non-redundant datasets contained a broad taxonomic representation and were used for downstream phylogenetic analyses.

#### Custom database of genomic resources

The custom database comprised genomic and transcriptomic resources from Porifera and representative opisthokont reference species (vertebrates*: Homo sapiens, Xenopus laevis, Danio rerio;* invertebrates*: Drosophila melanogaster, Caenorhabditis elegans, Nematostella vectensis, Mnemiopsis leidy;* fungi*: Aspergillus niger, Saccharomyces cerevisiae;* Table S8). For taxa lacking assembled transcriptomes, *de novo* transcriptome assemblies were generated using TRINITY v.2.15.1^78^, and protein-coding sequences were predicted using TRANSDECODER v.5.5.0 with default parameters.

#### Candidate validation and curation

For each *coq* gene family, recovered sequences were aligned and analyzed separately. Multiple sequence alignments were generated using MAFFT v7.490^53^ with default parameters. Maximum-likelihood phylogenetic inference using IQ-TREE v2.2.2.6^55,56^ under the best-fitting evolutionary model (Table S11) and 1000 ultrafast bootstraps. Phylogenetic trees were inspected manually to assess potential contamination. Sequences that branched robustly within bacterial clades or outside the expected Coq/Ubi family context were considered likely contaminants and were excluded from downstream analyses. Presence–absence patterns of *coq* genes across taxa were inferred based on the curated phylogenetic datasets.

### Tracing dietary UQ as a precursor for RQ biosynthesis in *Ephydatia muelleri*

To test whether sponges can convert exogenous UQ to RQ, we generated *Escherichia coli* producing ^13^C_6_-labelled UQ_8_. We then fed these bacteria to laboratory-reared adult *E. muelleri* and quantified UQ_8_ and RQ_8_ isotopologues using LC–MS.

#### Production of ^13^C_6_-labelled UQ_8_ in Escherichia coli

A *ubiC* knockout *E. coli* strain^79^ was cultured at 37 °C and 180 rpm shaking in M9 medium containing 0.4% glucose^80^. A 5 mL overnight preculture was used to inoculate a 50 mL preculture, which was grown overnight and subsequently scaled up to a 1 L culture. The culture was supplemented with 10 µM 13C₇-4-hydroxybenzoic acid (13C₇-4HB; Sigma-Aldrich) and incubated for 5 h. Cells were harvested by centrifugation (10,000 × *g*, 10 min, 4 °C), washed with 40 mL ice-cold PBS, pelleted (4,000 × *g*, 15 min, 4 °C), and stored at −80 °C. LC-MS analysis of a cell aliquot confirmed a 97.8% enrichment of the 13C₆ isotopologue of UQ_8_. Before feeding to sponges, bacterial cells were heat-killed at 70 °C for 30 min, and stored at −20 °C until further use.

#### Ephydatia muelleri exposure to ^13^C_6_-UQ_8_–labelled E. coli

Gemmules of the freshwater sponge *E. muelleri* (O’Connor Lake, Canada) were induced to hatch under laboratory conditions (Strekal’s Medium, 20 °C, constant darkness). At developmental stage 5, sponges were assigned to one of three treatments: (i) exposure to ^13^C_6_-UQ_8_–labelled *E. coli* (5 x 10^6^ cells/sponge), (ii) exposure to unlabeled *E. coli* (strain 5hda; 5 x 10^6^ cells/sponge), or (iii) no-food control. After 7 d of daily feeding, sponge biomass was harvested and placed at −80 °C until processing for quinone extraction.

#### Quantification of UQ_8_ and RQ_8_ isotopologues by LC–MS

UQ_8_ and RQ_8_ were quantified by LC–MS, and incorporation of the ^13^C_6_ label was assessed based on isotopologue-specific mass-to-charge (m/z) ratios. Rhodoquinone derived from utilization of the ^13^C_6_-UQ_8_–labelled *E. coli* was expected to show a predominant +6 Da m/z shift relative to rhodoquinone detected in sponges exposed to unlabeled *E. coli* (Table S9).

### Identification of electron transport chain components in sponges

Predicted proteomes of marine and freshwater sponges were screened for main subunits of the mitochondrial respiratory complexes (CI-CV). Encoded subunits from *Homo sapiens* were used as queries in similarity-based searches (blastp) against the sponge predicted proteomes (Supplementary Data 11). Protein family and taxonomic origin of each Porifera sequence were confirmed with a reciprocal blastp search against *nr* (Accessed March 2026) and using interproscan (v5.73-104.0).

### Identification and comparison of quinone-interacting residues in complex II

To examine sequence variation in the quinone-binding region of complex II, SdhC protein sequences were extracted from freshwater and marine sponge taxa, as well as selected metazoan reference species, and RquA bearing Opisthokonts. As there are no annotated gene models for the transcriptomes of *Paramoeba* and the genome of *Quaeritorhiza haematococci,* we predicted their putative *sdhC* coding sequences based on homology using tblastn (accessed March 2026). The *sdhC* gene sequence in *Quaeritorhiza haematococci* was identified and annotated through iterative homology-based alignments, with intron boundaries predicted using canonical GT-AG splice site motifs. To identify the conservation of amino acid sites in SdhC all metazoan sequences in the InterPro database matching HMM IPR014314 (SdhC) were retrieved and filtered by length (100-200 AA). These sequences were added to an in-house curated dataset of porifera, nematode, and Platyhelminthes sequences and aligned using MAFFT v7.490^53^ with the FFT-NS-2 algorithm. We identified Q-binding sites based on the positional homology to known Q-binding sites^37–39^ and numbered these sites from 1-5. From this dataset, we selected SdhC sequences from representative poriferan, opisthokonts (including *Q. haematococci*), and *Paramoeba* species) and refined the alignment using MAFFT L-INS-i v7.490. Alignments were manually inspected to identify conserved and variable positions in sites implicated in quinone interaction.

### Structural modeling and *in silico* docking analyses

SDH proteins for Homo sapiens were retrieved from UniProt (SDHA: P31040; SDHB: P21912, SDHC: Q99643, SDHD: O14521). SDH proteins from E. muelleri were retrieved using blastp against GCA_013339895.126, and when gene models were not predicted, we queried the transcriptome using tblastn using the human sequences as queries. Mitochondrial transit peptides were removed from sequences using TargetP 2.0^81^. Multple sequence alignments (MSAs) for each chain were generated via the AlphaFold3 web server, and the resulting paired MSAs were used as input for structure prediction of each complex^82^. Rhodoquinone-10 (RQ_10_), Rhodoquinol-10 (RQH_10_), Rhodoquinone-1 (RQ_1_), Rhodoquinol-1 (RQH_1_), Ubiquinone-10 (UQ_10_), Ubiquinol-10 (UQH_10_), Ubiquinone-1 (UQ_1_) and Ubiquinol-1 (UQH_1_) were each docked in CII sequences using Boltz-2 supplied with the AlphaFold3 paired alignments^83^, with Cofactors flavin-adenine dinucleotide and protoporphyrin IX (without Fe) and lipid l-alpha-phosphatidyl-beta-oleoyl-gamma-palmitoyl-phosphatidylethanolamine (PE) included in all predictions. A contact constraint of 4 Å between the quinone and the conserved serine (Fig S6, site 5) was implemented in Boltz-2 (with a molecular weight correction to normalize across different sized ligands). Docking simulations were repeated across five independent runs to account for stochastic variation in ligand scoring, and the first affinity prediction value (negative log10(IC50), where IC50 (half maximal inhibitory concentration) is in μM, was taken from each run and converted to ΔG (ΔG [kcal/mol] ≈ -1.364 × (6 - prediction value^83^. Docking affinity (ΔG) values were compared across quinone ligands separately for each species (*E. muelleri* and *H. sapiens*) using Welch’s one-way ANOVA. *Post hoc* comparisons were performed using the Games-Howell correction. Statistical significance was set at a p-adjusted < 0.05.

## Acknowledgements

We thank Gordon Lax for providing us with Euglena RquA sequences, Malcolm Hill for assistance with marine sponge sample collection, and Kaan Dizer for helping us with sponge rearing. We also thank Ola Gustafsson and Mark Lessard for helping us with confocal imaging. We also thank Sophie Abby for valuable discussions and feedback during the experimental process. Computational resources and data handling were enabled by the National Academic Infrastructure for Supercomputing in Sweden (NAISS), partially funded by the Swedish Research Council through grant agreement no. 2022 -06725, under projects NAISS 2024/5-77, NAISS 2024/6-43, NAISS 2025/22-280 and NAISS 2025/5-253 with access to the UPPMAX and PDC computational infrastructures. We also thank Victor Törnblom for maintaining our local server used for some of the analyses. This project was partially funded from the Carl Tryggers Stiftelsen (CTS 21:1693 awarded to CWS), vetenskapsrådet starting grant (2020-05071 awarded to CWS), and Kungliga Fysiografiska Sällskapet i Lund (44216, 45410, 43424, awarded to SP). Research in the AS-P group was supported by the European Research Council (ERC-StG 851647) and the Spanish Ministry of Science and Innovation (PID2021-124757NB-I00). Research in Shepherd Lab was supported by Scholl Distinguished Professor Award (J.N.S.), Gonzaga Science Research Program (P.E.C.), and Kay Nakamaye Research Award (L.N.B.).

## Author contributions

Field Sample collection: SP, RP, ALH, SPL

Data collection: SP, MV, NB, PEC, HI, RP, TNA, LNB, KIAC, LF, DS, CSW, FP, JNS, CWS

Data analyses: SP, MV, NB, PEC, JB, RP, IVK, LNB, AS-B, JNS, ALH, CWS

Visualization: SP, NB, JB, IVK, HI, KIAC, CWS

Interpretation: SP, MV, NB, JB, FP, JNS, ALH, SPL, CWS

Writing: Drafted by SP incorporating materials from JB, MV, NB, HI, IVK, FP, JNS, and CWS. All authors revised and approved the final version.

## Data and Code availability

All data and code are available here: https://figshare.com/s/fc115a87d18730521ca3 and will become publicly available upon publication with the DOI 10.6084/m9.figshare.31463641.

## References

1. Ochman, H., Lawrence, J. G. & Groisman, E. A. Lateral gene transfer and the nature of bacterial innovation. Nature 405, 299–304 (2000).

2. Eme, L., Gentekaki, E., Curtis, B., Archibald, J. M. & Roger, A. J. Lateral Gene Transfer in the Adaptation of the Anaerobic Parasite Blastocystis to the Gut. Curr. Biol. 27, 807–820 (2017).

3. Stairs, C. W. et al. Microbial eukaryotes have adapted to hypoxia by horizontal acquisitions of a gene involved in rhodoquinone biosynthesis. eLife 7, e34292 (2018).

4. Stairs, C. W., Leger, M. M. & Roger, A. J. Diversity and origins of anaerobic metabolism in mitochondria and related organelles. Philos. Trans. R. Soc. B Biol. Sci. 370, 20140326 (2015).

5. Andersson, J. O. Lateral gene transfer in eukaryotes. Cell. Mol. Life Sci. 62, 1182–1197 (2005).

6. Ku, C. & Martin, W. F. A natural barrier to lateral gene transfer from prokaryotes to eukaryotes revealed from genomes: the 70 % rule. BMC Biol. 14, 89 (2016).

7. Martin, W. F. Too Much Eukaryote LGT. BioEssays 39, 1700115 (2017).

8. Keeling, P. J. Horizontal gene transfer in eukaryotes: aligning theory with data. Nat. Rev. Genet. 25, 416–430 (2024).

9. Danchin, E. G. J., Guzeeva, E. A., Mantelin, S., Berepiki, A. & Jones, J. T. Horizontal Gene Transfer from Bacteria Has Enabled the Plant-Parasitic Nematode *Globodera pallida* to Feed on Host-Derived Sucrose. Mol. Biol. Evol. 33, 1571–1579 (2016).

10. Nováková, E. & Moran, N. A. Diversification of Genes for Carotenoid Biosynthesis in Aphids following an Ancient Transfer from a Fungus. Mol. Biol. Evol. 29, 313–323 (2012).

11. Nowell, R. W. et al. Bdelloid rotifers deploy horizontally acquired biosynthetic genes against a fungal pathogen. Nat. Commun. 15, 5787 (2024).

12. Raymond, J. A. A horizontally transferred bacterial gene aids the freezing tolerance of Antarctic bdelloid rotifers. Proc. Natl. Acad. Sci. 122, e2421910122 (2025).

13. Gawryluk, R. M. R. & Stairs, C. W. Diversity of electron transport chains in anaerobic protists. Biochim. Biophys. Acta BBA - Bioenerg. 1862, 148334 (2021).

14. Salinas, G., Langelaan, D. N. & Shepherd, J. N. Rhodoquinone in bacteria and animals: Two distinct pathways for biosynthesis of this key electron transporter used in anaerobic bioenergetics. Biochim. Biophys. Acta BBA - Bioenerg. 1861, 148278 (2020).

15. Guerra, R. M. & Pagliarini, D. J. Coenzyme Q biochemistry and biosynthesis. Trends Biochem. Sci. 48, 463–476 (2023).

16. Stefely, J. A. & Pagliarini, D. J. Biochemistry of Mitochondrial Coenzyme Q Biosynthesis. Trends Biochem. Sci. 42, 824–843 (2017).

17. Roberts Buceta, P. M., et al. The kynurenine pathway is essential for rhodoquinone biosynthesis in Caenorhabditis elegans. J. Biol. Chem. 294, 11047–11053 (2019).

18. Tan, J. H. et al. Alternative splicing of coq-2 controls the levels of rhodoquinone in animals. eLife 9, e56376 (2020).

19. Lonjers, Z. T. et al. Identification of a New Gene Required for the Biosynthesis of Rhodoquinone in Rhodospirillum rubrum. J. Bacteriol. 194, 965–971 (2012).

20. Simion, P. et al. A Large and Consistent Phylogenomic Dataset Supports Sponges as the Sister Group to All Other Animals. Curr. Biol. 27, 958–967 (2017).

21. Nettersheim, B. J. et al. Putative sponge biomarkers in unicellular Rhizaria question an early rise of animals. *Nat*. Ecol. Evol. 3, 577–581 (2019).

22. Mentel, M., Röttger, M., Leys, S., Tielens, A. G. M. & Martin, W. F. Of early animals, anaerobic mitochondria, and a modern sponge. BioEssays 36, 924–932 (2014).

23. Mills, D. B. et al. Oxygen requirements of the earliest animals. Proc. Natl. Acad. Sci. 111, 4168–4172 (2014).

24. Mills, D. B. & Sperling, E. A. Marine sponges in the once and future ocean. Glob. Change Biol. 28, 1953–1955 (2022).

25. Micaroni, V. et al. Adaptive strategies of sponges to deoxygenated oceans. Glob. Change Biol. 28, 1972–1989 (2022).

26. Kenny, N. J. et al. Tracing animal genomic evolution with the chromosomal-level assembly of the freshwater sponge Ephydatia muelleri. Nat. Commun. 11, 3676 (2020).

27. Leys, S. P. et al. A morphological cell atlas of the freshwater sponge Ephydatia muelleri with key insights from targeted single-cell transcriptomes. EvoDevo 16, 1 (2025).

28. Hernández Márquez, S., et al. The genus *Eutreptiella* (Euglenophyceae/Euglenozoa) across its global distribution range. Ecol. Evol. 14, e70241 (2024).

29. Hoffmeister, M. et al. Euglena gracilis Rhodoquinone:Ubiquinone Ratio and Mitochondrial Proteome Differ under Aerobic and Anaerobic Conditions. J. Biol. Chem. 279, 22422–22429 (2004).

30. Burki, F., Roger, A. J., Brown, M. W. & Simpson, A. G. B. The New Tree of Eukaryotes. Trends Ecol. Evol. 35, 43–55 (2020).

31. Bernert, A. C. et al. Recombinant RquA catalyzes the in vivo conversion of ubiquinone to rhodoquinone in Escherichia coli and Saccharomyces cerevisiae. Biochim. Biophys. Acta BBA - Mol. Cell Biol. Lipids 1864, 1226–1234 (2019).

32. Pfaffl, M. W. A new mathematical model for relative quantification in real-time RT-PCR. Nucleic Acids Res. 29, 45e–445 (2001).

33. Neupane, T. et al. Microbial rhodoquinone biosynthesis proceeds via an atypical RquA-catalyzed amino transfer from S-adenosyl-L-methionine to ubiquinone. Commun. Chem. 5, 89 (2022).

34. Pelosi, L. et al. COQ4 is required for the oxidative decarboxylation of the C1 carbon of coenzyme Q in eukaryotic cells. Mol. Cell 84, 981–989.e7 (2024).

35. Maklashina, E. & Cecchini, G. The quinone-binding and catalytic site of complex II. Biochim. Biophys. Acta BBA - Bioenerg. 1797, 1877–1882 (2010).

36. Horsefield, R. et al. Structural and Computational Analysis of the Quinone-binding Site of Complex II (Succinate-Ubiquinone Oxidoreductase). J. Biol. Chem. 281, 7309–7316 (2006).

37. Sun, F. et al. Crystal Structure of Mitochondrial Respiratory Membrane Protein Complex II. Cell 121, 1043–57 (2005).

38. Du, Z. et al. Structure of the human respiratory complex II. Proc. Natl. Acad. Sci. 120, e2216713120 (2023).

39. Inaoka, D. et al. Structural Insights into the Molecular Design of Flutolanil Derivatives Targeted for Fumarate Respiration of Parasite Mitochondria. Int. J. Mol. Sci. 16, 15287–15308 (2015).

40. Molčanov, K. & Kojić-Prodić, B. Towards understanding π-stacking interactions between non-aromatic rings. IUCrJ 6, 156–166 (2019).

41. Holcomb, M., Chang, Y., Goodsell, D. S. & Forli, S. Evaluation of ALPHAFOLD2 structures as docking targets. Protein Sci. 32, e4530 (2023).

42. Wetzel, R. G. Limnology: Lake and River Ecosystems. (Elsevier Academic Press, 2001).

43. Valeros, J. et al. Rhodoquinone carries electrons in the mammalian electron transport chain. Cell 188, 1084–1099.e27 (2025).

44. Simpson, T. L. The Cell Biology of Sponges. (1984).

45. Imsiecke, G. Ingestion, digestion, and egestion in Spongilla lacustris (Porifera, Spongillidae) after pulse feeding with Chlamydomonas reinhardtii (Volvocales). Zoomorphology 113, 233–244 (1993).

46. Leys, S. P. & Hill, A. The Physiology and Molecular Biology of Sponge Tissues. In Advances in Marine Biology vol. 62 1–56 (Elsevier, 2012).

47. Jensen, L., Grant, J. R., Laughinghouse, H. D. & Katz, L. A. Assessing the effects of a sequestered germline on interdomain lateral gene transfer in Metazoa. Evolution 70, 1322–1333 (2016).

48. Kumala, L., Larsen, M., Glud, R. N. & Canfield, D. E. Spatial and temporal anoxia in single-osculum *Halichondria panicea* demosponge explants studied with planar optodes. Mar. Biol. 168, 173 (2021).

49. Reiswig, H. M. & Miller, T. L. Freshwater Sponge Gemmules Survive Months of Anoxia. Invertebr. Biol. 117, 1 (1998).

50. Sokolova, A. M. & Ereskovsky, A. V. How gemmules become sponges: known facts and open questions. Invertebr. Zool. 22, 383–400 (2025).

51. Richter, D. J., et al. EukProt: A database of genome-scale predicted proteins across the diversity of eukaryotes. Peer Community J. 2, e56 (2022).

52. Lax, G. & Simpson, A. G. B. The molecular diversity of phagotrophic Euglenids examined using single-cell methods. 10.5061/dryad.m0cfxpp0s.

53. Katoh, K. MAFFT: a novel method for rapid multiple sequence alignment based on fast Fourier transform. Nucleic Acids Res. 30, 3059–3066 (2002).

54. Criscuolo, A. & Gribaldo, S. BMGE (Block Mapping and Gathering with Entropy): a new software for selection of phylogenetic informative regions from multiple sequence alignments. BMC Evol. Biol. 10, 210 (2010).

55. Minh, B. Q. et al. IQ-TREE 2: New Models and Efficient Methods for Phylogenetic Inference in the Genomic Era. Mol. Biol. Evol. 37, 1530–1534 (2020).

56. Hoang, D. T., Chernomor, O., Von Haeseler, A., Minh, B. Q. & Vinh, L. S. UFBoot2: Improving the Ultrafast Bootstrap Approximation. Mol. Biol. Evol. 35, 518–522 (2018).

57. Wang, H. C., Minh, B. Q., Susko, E. & Roger, A. J. Modeling Site Heterogeneity with Posterior Mean Site Frequency Profiles Accelerates Accurate Phylogenomic Estimation. 67, 216–235 (2018).

58. Lemoine, F. et al. Renewing Felsenstein’s phylogenetic bootstrap in the era of big data. Nature 556, 452–456 (2018).

59. Felsenstein, J. Confedence limits on phylogenies: an approach using the bootstrap. 39, 783–791.

60. Kim, I. V. et al. Chromatin loops are an ancestral hallmark of the animal regulatory genome. Nature 642, 1097–1105 (2025).

61. Open2C et al. Pairtools: From sequencing data to chromosome contacts. PLOS Comput. Biol. 20, e1012164 (2024).

62. Abdennur, N. & Mirny, L. A. Cooler: scalable storage for Hi-C data and other genomically labeled arrays. Bioinformatics 36, 311–316 (2020).

63. Xu, W. et al. CoolBox: a flexible toolkit for visual analysis of genomics data. BMC Bioinformatics 22, 489 (2021).

64. Dobin, A. et al. STAR: ultrafast universal RNA-seq aligner. Bioinformatics 29, 15–21 (2013).

65. Ramírez, F. et al. deepTools2: a next generation web server for deep-sequencing data analysis. Nucleic Acids Res. 44, W160–W165 (2016).

66. Quinlan, A. R. & Hall, I. M. BEDTools: a flexible suite of utilities for comparing genomic features. Bioinformatics 26, 841–842 (2010).

67. Bailey, T. L., Johnson, J., Grant, C. E. & Noble, W. S. The MEME Suite. Nucleic Acids Res. 43, W39–W49 (2015).

68. Grant, C. E., Bailey, T. L. & Noble, W. S. FIMO: scanning for occurrences of a given motif. Bioinformatics 27, 1017–1018 (2011).

69. Gupta, S., Stamatoyannopoulos, J. A., Bailey, T. L. & Noble, W. S. Quantifying similarity between motifs. Genome Biol. 8, R24 (2007).

70. Ovek Baydar, D., et al. JASPAR 2026: expansion of transcription factor binding profiles and integration of deep learning models. Nucleic Acids Res. 54, D184–D193 (2026).

71. Gardiner-Garden, M. & Frommer, M. CpG Islands in vertebrate genomes. J. Mol. Biol. 196, 261–282 (1987).

72. Parker, J. L. & Newstead, S. Method to increase the yield of eukaryotic membrane protein expression in *SACCHAROMYCES CEREVISIAE* for structural and functional studies. Protein Sci. 23, 1309–1314 (2014).

73. Robert, C., Pereira, R. & Thollesson, M. Addition to Sweden’s freshwater sponge fauna and a phylogeographic study of Spongilla lacustris (Spongillida, Porifera) in southern Sweden. Eur. J. Taxon. 828, (2022).

74. Riesgo, A. et al. Transcriptomic analysis of differential host gene expression upon uptake of symbionts: a case study with Symbiodinium and the major bioeroding sponge Cliona varians. BMC Genomics 15, 376 (2014).

75. Schindelin, J., et al. Fiji: an open-source platform for biological-image analysis. Nat. Methods 9, 676–682 (2012).

76. Ye, J. et al. Primer-BLAST: A tool to design target-specific primers for polymerase chain reaction. BMC Bioinformatics 13, 134 (2012).

77. Ramakers, C., Ruijter, J. M., Deprez, R. H. L. & Moorman, A. F. M. Assumption-free analysis of quantitative real-time polymerase chain reaction (PCR) data. Neurosci. Lett. 339, 62–66 (2003).

78. Grabherr, M. G. et al. Full-length transcriptome assembly from RNA-Seq data without a reference genome. Nat. Biotechnol. 29, 644–652 (2011).

79. Pelosi, L. et al. Ubiquinone Biosynthesis over the Entire O2 Range: Characterization of a Conserved O2-Independent Pathway. mBio 10, e:01319-19 (2019).

80. Roger-Margueritat, M. et al. Heterologous plastoquinone production using a newly identified O_2_ -dependent cyanobacterial hydroxylase. FEBS J. 293, 1708–1726 (2026).

81. Almagro Armenteros, J. J., et al. Detecting sequence signals in targeting peptides using deep learning. Life Sci. Alliance 2, e201900429 (2019).

82. Abramson, J. et al. Accurate structure prediction of biomolecular interactions with AlphaFold 3. Nature 630, 493–500 (2024).

83. Boltz-2: Towards Accurate and Efficient Binding Affinity Prediction. *bioRxiv* doi: 10.1101/2025.06.14.659707 (2025).

