## Supplementary Figures S1-S7 for "Lateral gene transfer introduced the microbial anaerobiosis-related gene *rquA* into early animals"

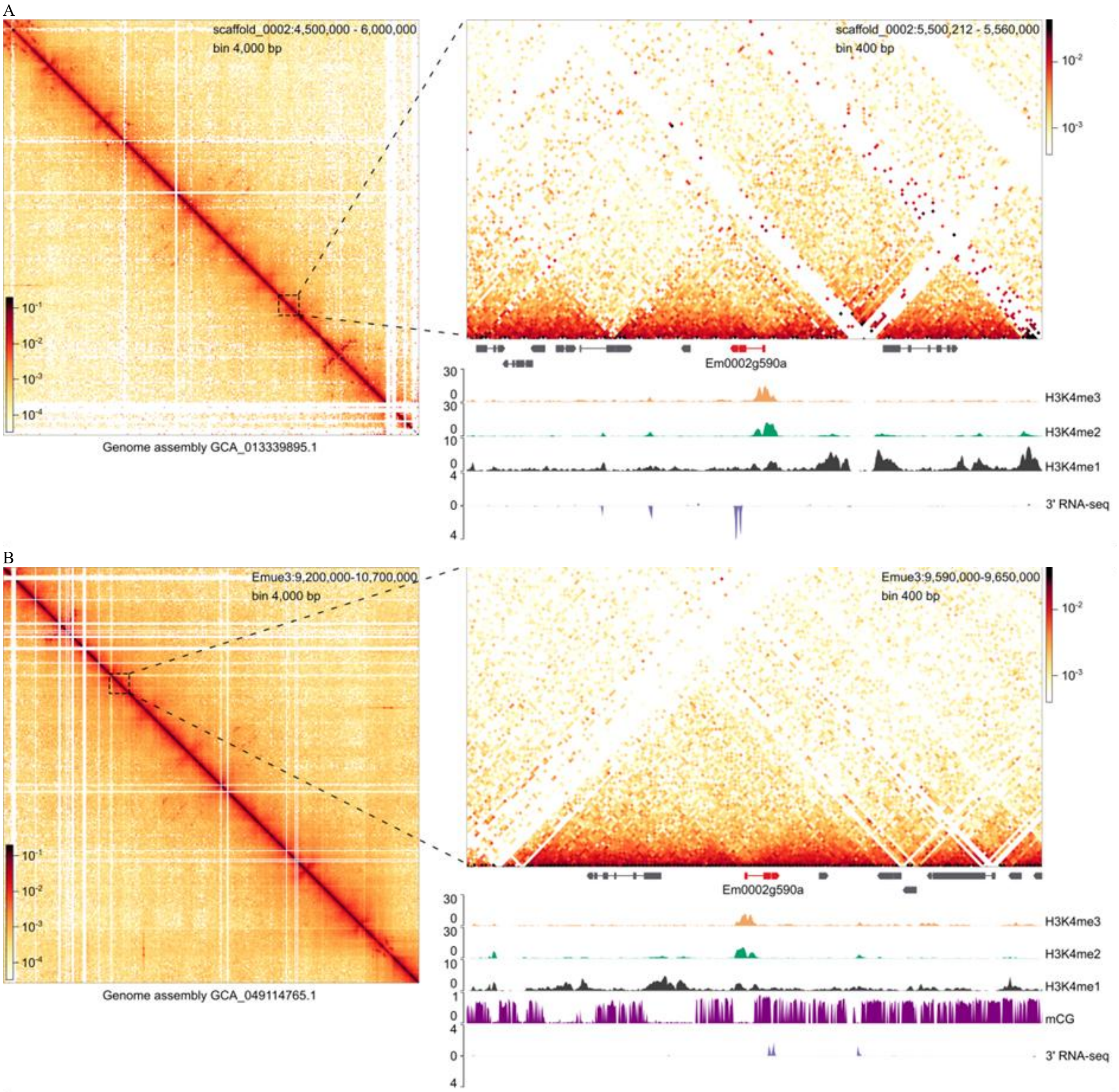


**Fig. S1. Local chromatin context of *Em0002g590a* in *Ephydatia muelleri*. A,** Chromatin contact map showing local *cis* interaction frequencies across the genomic locus harboring *Em0002g590a* gene (*rquA*). *Em0002g590a* (red) has a continuous contact pattern without an observable disruption in interaction intensity at the locus. Tracks display histone modifications (H3K4me3, H3K4me2, H3K4me1), CpG methylation (mCG), and transcript abundance in choanocytes (3′ RNA-seq) across the region. Enrichment of H3K4me3 signal is observed at the promoter of *Em0002g590a*. Interaction maps were balanced using iterative correction and eigenvector decomposition (ICE) normalization[1]. **B,** Same as (A) but for contact data aligned to the reference genome assembly GCA_049114765.1.

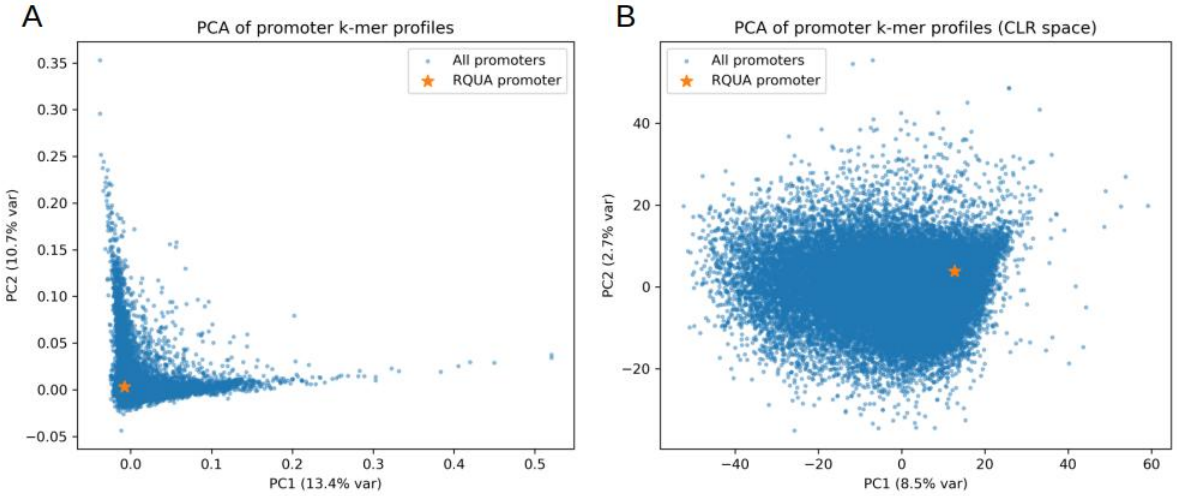


**Fig. S2: Principal component analysis (PCA) of 4-mer promoter composition. A,** PCA was performed on centered relative frequencies of all 4-mers derived from promoter sequences. Each point represents one promoter k-mer profile. **B,** CLR-transformed PCA (CLR-PCA) of promoter 4-mer profiles. Because k-mer frequencies are compositional, a centered log-ratio (CLR) transform was applied prior to PCA to remove closure effects. In both panels, the *rquA* promoter is highlighted as an orange star.

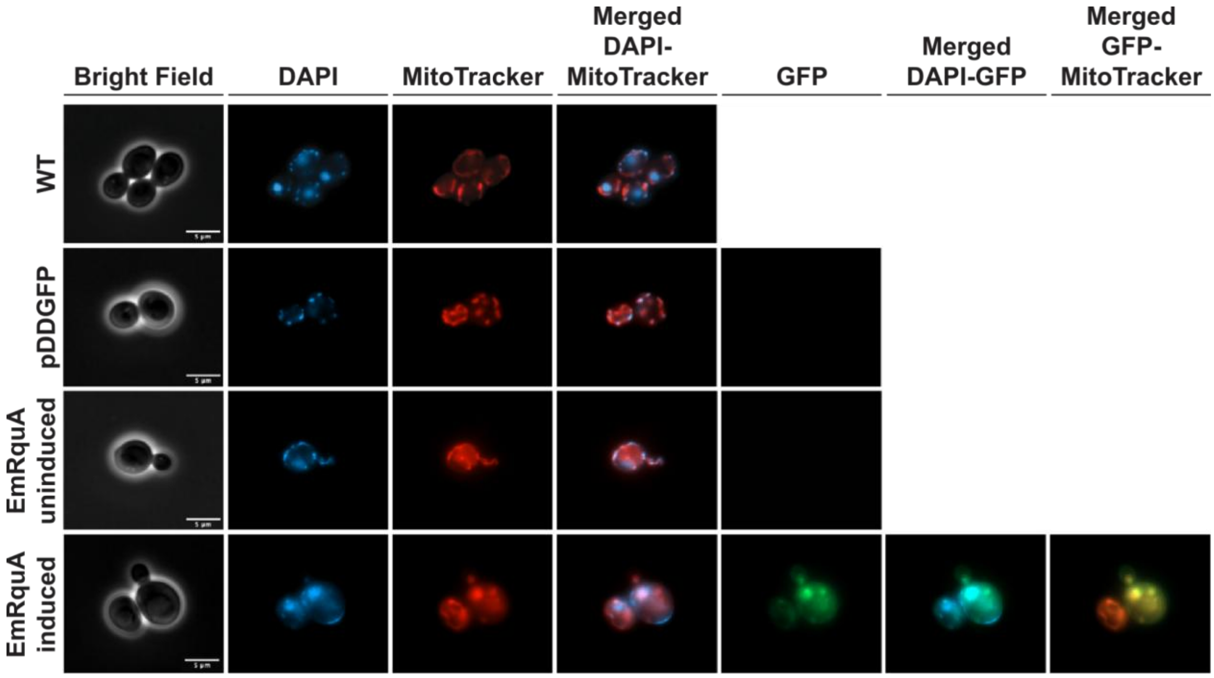


**Fig. S3: Mitochondrial localization of sponge Em-rquA in yeast.** Representative fluorescence microscopy images of *Saccharomyces cerevisiae* expressing GFP-tagged *Ephydatia muelleri* rquA (*Em*-RquA, induced) and corresponding controls. From left to right, panels show brightfield, DNA staining (DAPI, blue), mitochondrial staining (MitoTracker, red), merged DAPI/MitoTracker, GFP fluorescence (GFP, green), and merged channels. W303 wild-type (WT) cells were grown in glucose (shown) and galactose (not shown). Cells containing the empty-vector (pDDGFP) were grown in galactose (expression inducing media) and had no GFP signal detected. In *Em*-RquA–expressing cells (induced, grown in galactose), the GFP signal overlapped with mitochondrial staining, whereas no mitochondrial GFP signal was observed in uninduced cells (grown in glucose). Scale bars = 5 µm.

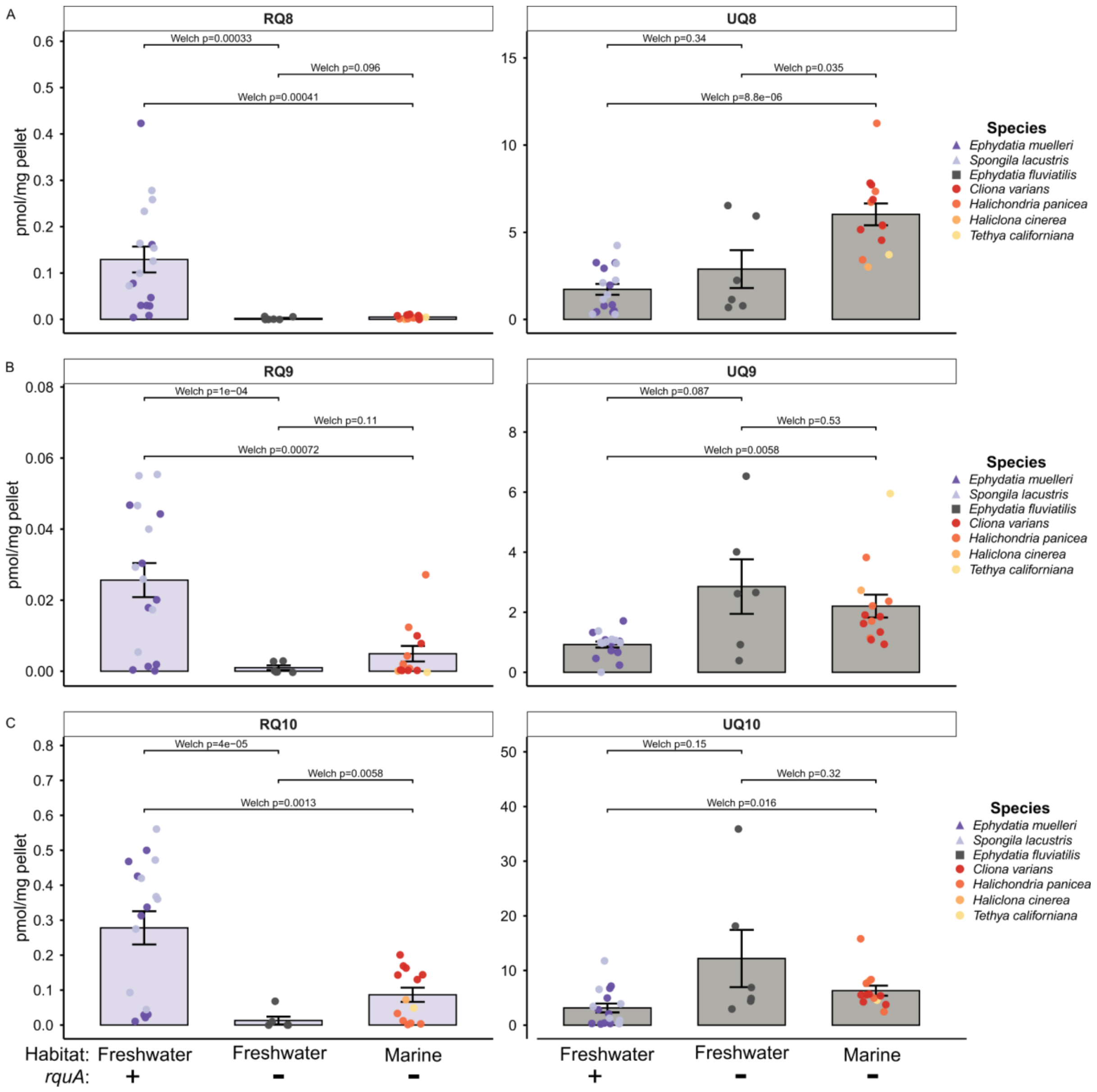


**Fig. S4. Quinone abundances across freshwater and marine sponges.** Absolute abundances of rhodoquinone ($RQ_8$, $RQ_9$, $RQ_{10}$; left panels) and ubiquinone ($UQ_8$, $UQ_9$, $UQ_{10}$; right panels) measured by LC-MS across different sponge species. Each data point represents an individual biological sample, and bar plots indicate group means with error bars showing ± s.e. Bottom labels indicate the presence (+) or absence (–) of RquA.

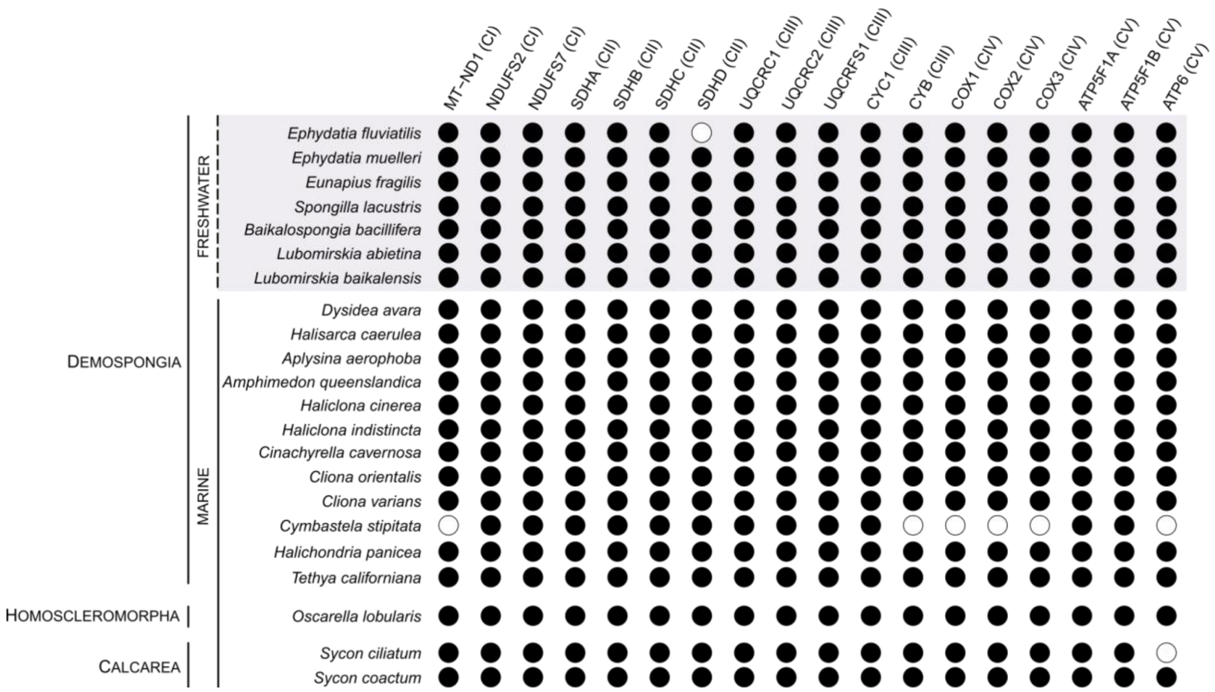


**Fig. S5. Presence and absence of mitochondrial electron transport chain (ETC) subunits across sponge species.** Rows represent species, and columns represent main subunits of ETC complexes CI–CV. Filled circles indicate presence of orthologs based on protein sequence analysis, whereas open circles indicate absence. Complex assignments are indicated in parentheses for each subunit.

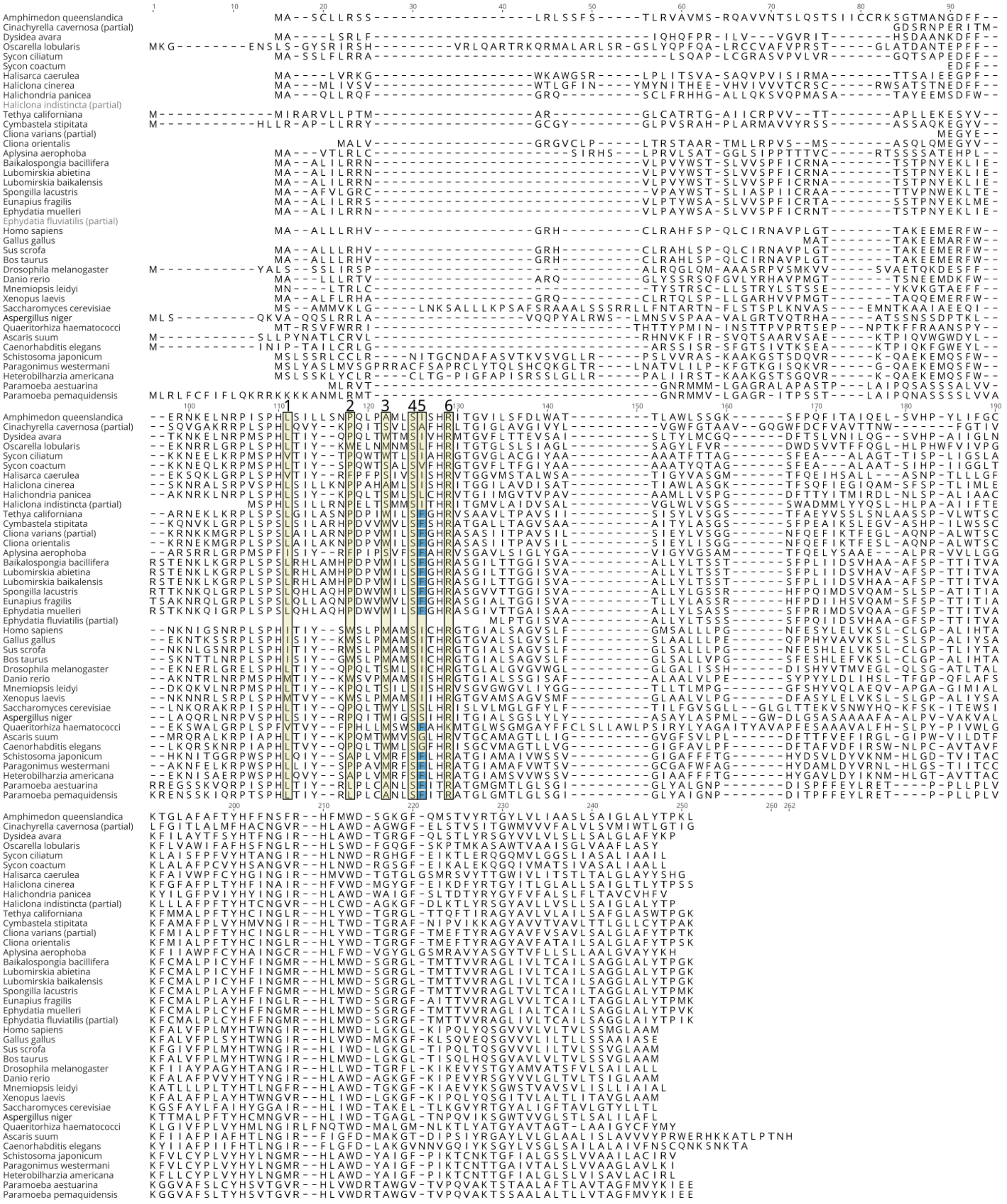

**Fig. S6. Alignment of SdhC protein sequences across sponge taxa, *rquA*-encoding, and RQ-producing species.** Numbered positions indicate quinone-binding sites identified in structural studies of different model organisms (*Ascaris suum*: sites 4, 5, and 6; *Homo sapiens*: sites 1, 2, 3, 5, and 6; *Sus scrofa*: sites 4 and 6)[2-4].

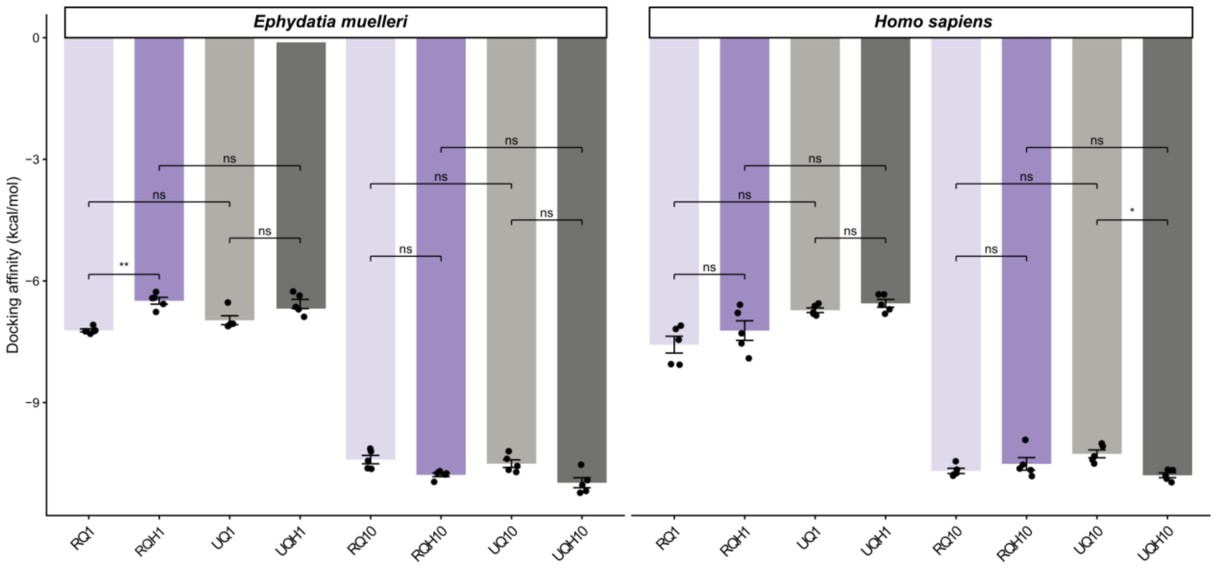


**Fig. S7. Docking affinities (kcal/mol) of RQ and UQ ligands in *E. muelleri* and *H. sapiens*.** Each dot represents a docking simulation with Boltz-2 and with distinct stochastic starting conditions. Statistical comparisons indicate no significant difference in binding between the two quinone ligands (UQ vs RQ) within either species. P-values represent adjusted p-values corrected with BH for multiple testing.

### References


1. Open2C; Abdennur, N. *et al*., Cooltools: Enabling high-resolution Hi-C analysis in Python. *PLoS Comput. Biol.* **20**, e1012067 (2024).
2. Inaoka, D. K. *et al.* Structural Insights into the Molecular Design of Flutolanil Derivatives Targeted for Fumarate Respiration of Parasite Mitochondria. *Int. J. Mol. Sci.* ***16***, 15287-15308 (2015).
3. Du, Z. *et al.* Structure of the human respiratory complex II, *Proc. Natl. Acad. Sci.* **120**, e2216713120 (2023).
4. Sun, F. *et al.* Crystal Structure of Mitochondrial Respiratory Membrane Protein Complex II, *Cell* **121**, 1043-1057 (2005).
